# MdfA is a novel ClpC adaptor protein that functions in the developing *Bacillus subtilis* spore

**DOI:** 10.1101/2024.03.02.583065

**Authors:** Shawn C. Massoni, Nicola J. Evans, Ingo Hantke, Colleen Fenton, James H. Torpey, Katherine M. Collins, Ewelina M. Krysztofinska, Janina H. Muench, Arjun Thapaliya, Santiago Martínez-Lumbreras, Sé Hart Martin, Celia Slater, Xinyue Wang, Ruth Fekade, Sandra Obwar, Siyu Yin, Alishba Vazquez, Christopher B. Prior, Kürşad Turgay, Rivka L. Isaacson, Amy H. Camp

## Abstract

Bacterial protein degradation machinery consists of chaperone-protease complexes that play vital roles in bacterial growth and development and have sparked interest as novel antimicrobial targets. ClpC-ClpP (ClpCP) is one such chaperone-protease, recruited by adaptors to specific functions in the model bacterium *Bacillus subtilis* and other Gram-positive bacteria including the pathogens *Staphylococcus aureus* and *Mycobacterium tuberculosis*. Here we have identified a new ClpCP adaptor protein, MdfA (“Metabolic differentiation factor A”, formerly YjbA), in a genetic screen for factors that help drive *B. subtilis* toward metabolic dormancy during spore formation. A knockout of *mdfA* stimulates gene expression in the developing spore, while aberrant expression of *mdfA* during vegetative growth is toxic. MdfA binds directly to ClpC to induce its oligomerization and ATPase activity, and this interaction is required for the in vivo effects of *mdfA*. Finally, a co-crystal structure reveals that MdfA binds to the ClpC N-terminal domain at a location analogous to that on the *M. tuberculosis* ClpC1 protein where bactericidal cyclic peptides bind. Altogether our data and that of an accompanying study by Riley, Lyda and colleagues support a model in which MdfA induces ClpCP-mediated degradation of metabolic enzymes in the developing spore, helping drive it toward metabolic dormancy.

## INTRODUCTION

Cells across all domains of life employ AAA+ proteases to degrade misfolded or aggregated proteins, as well as to control the abundance of specific proteins and, in turn, the myriad biological processes they regulate (Sauer and Baker 2011). The most complex and perhaps best-known AAA+ protease, the 26S proteasome, is responsible for the majority of protein degradation in eukaryotic cells, regulating essential processes including the cell cycle, signal transduction, and apoptosis (Collins and Goldberg 2017). In bacteria, the degradation of proteins by a more diverse set of AAA+ proteases (e.g. Clp, Lon, FtsH proteases) drives cell division, biofilm formation, pathogenesis, competence, and other developmental processes (Mahmoud and Chien 2018). Given their far-reaching roles in cell physiology, AAA+ proteases are attractive therapeutic targets. For example, proteasome inhibitors have been used to treat certain human cancers (Fricker 2020), and natural products that dysregulate bacterial AAA+ proteases can act as antibiotics against pathogens such as *Mycobacterium tuberculosis* and *Staphylococcus aureus* (Culp and Wright 2017).

Like the proteasome, bacterial AAA+ proteases are large barrel-like molecular machines with both chaperone and protease domains (Mahmoud and Chien 2018). The chaperone, a member of the Hsp100/Clp AAA+ (ATPases associated with various cellular activities) family, forms a hexamer and uses conformational changes induced by the binding and hydrolysis of ATP to unfold and translocate protein substrates through its central pore and into the interior chamber of the protease, where the proteolytic active sites are buried. Unlike the proteasome, which recognizes substrate proteins marked by the covalent addition of ubiquitin, bacterial AAA+ proteases typically recognize their substrates via intrinsic peptide sequences (“degrons”) to which the chaperone component binds directly or indirectly via adaptor proteins (Kuhlmann and Chien 2017). Adaptors can also bind to and influence the activity of a particular bacterial AAA+ proteases in other ways, for example by promoting assembly of an active complex (Kuhlmann and Chien 2017).

In Gram positive bacteria including the model bacterium *Bacillus subtilis*, the ClpC-ClpP (ClpCP) protease composed of the AAA+ chaperone ClpC and the protease ClpP plays a central role in general and regulatory proteolysis, impacting cellular pathways including stress resistance, competence, sporulation, and virulence (e.g. Gunaratnam et al. 2019; Krüger et al. 1994; Msadek et al. 1994; Rouquette et al. 1998; Pan et al. 2001; Meeske et al. 2016). In *M. tuberculosis*, the equivalent protease (ClpC1P1P2) is essential and the target of bactericidal cyclic peptides cyclomarin A, ecumicin, lassomycin, and rufomycin (Choules et al. 2019; Gao et al. 2015; Gavrish et al. 2014; Schmitt et al. 2011).

ClpC harbors two AAA+ ATPase domains, D1 and D2, that form stacked rings in the active hexamer (Sauer and Baker 2011; Wang et al. 2011). Loops within D2 allow the ClpC hexamer to dock onto two stacked heptameric rings of the ClpP protease. ClpC additionally contains a coiled-coil middle domain (M-domain) embedded within D1, as well as an N-terminal domain (N-domain) that sits atop the D1 ring and mediates selection of substrates for degradation. In some cases, the N-domain directly interacts with substrates, for example those that are marked with a phosphorylated arginine (pArg) residue (Trentini et al. 2016). In other cases, the ClpC N-domain indirectly interacts with substrates via adaptor proteins such as MecA, which recruits the *B. subtilis* competence master regulator ComK and other substrates for ClpCP-mediated degradation (Turgay et al. 1998).

Here we report the identification and characterization of a novel ClpCP adaptor protein that functions during *B. subtilis* endospore formation (“sporulation”). Sporulation is triggered by starvation and culminates in the production of an environmentally resistant, metabolically dormant spore (**Fig 1A**) (Tan and Ramamurthi 2014; Riley et al. 2020). This developmental process involves two cells that arise from an early asymmetric division event: a smaller “forespore” that becomes the spore, and a larger “mother cell” that aids in forespore development but ultimately dies. Soon after asymmetric division, while the two cells lie side-by-side, developmental gene expression is controlled by the alternative sigma (σ) factors σ^F^ in the forespore and σ^E^ in the mother cell. The mother cell then engulfs the forespore, giving rise to a cell-within-a-cell configuration, at which point σ^G^ and σ^K^ take over to drive late gene expression in the forespore and mother cell, respectively. In the final stages of development, the forespore acquires protective peptidoglycan cortex and protein coat layers, and upon lysis of the mother cell, is released into the environment as a mature spore.

**Figure 1.**
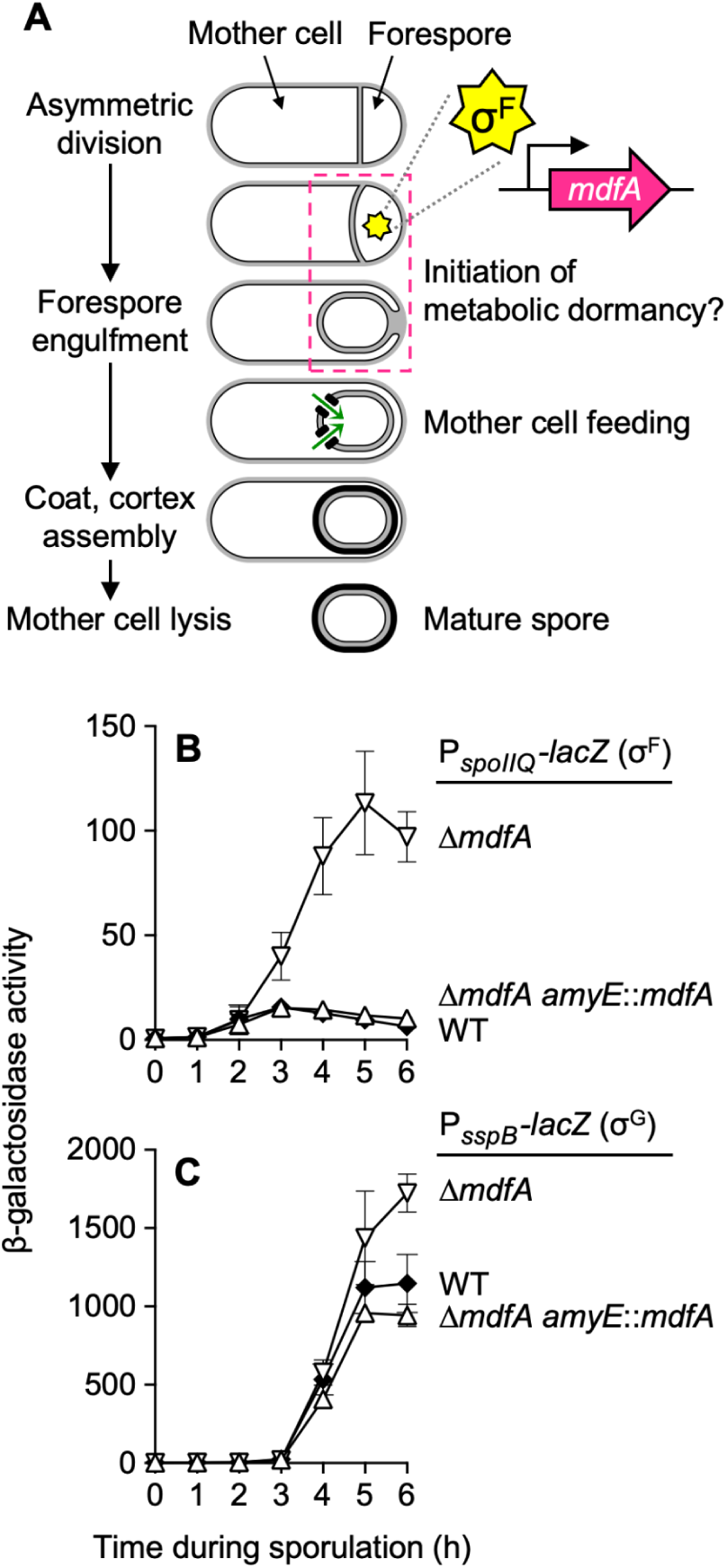
Forespore gene expression is stimulated in cells deleted for *mdfA*. **(A)** Cartoon representation of *B. subtilis* sporulation, with major morphological events indicated. Also shown is the SpoIIQ•SpoIIIAA-AH channel apparatus through which the mother cell provides small molecule metabolites to support the forespore during late development (Camp and Losick 2009; Riley et al. 2021). The early-acting forespore sigma factor σ^F^ is shown, as is the *mdfA* (formerly *yjbA*) gene, which has been assigned to the σ^F^ regulon (Arrieta-Ortiz et al. 2015; Steil et al. 2005; Wang et al. 2006) and is the focus of this study. **(B,C)** The forespore-specific sigma factors σ^F^ and σ^G^ are significantly more active in *ΔmdfA* cells. Production of β-galactosidase from the **(B)** σ^F^-dependent P*_spoIIQ_-lacZ* reporter or **(C)** σ^G^-dependent P*_sspB_-lacZ* reporter was monitored during sporulation of otherwise wild type cells (WT; closed diamonds), cells deleted for *mdfA* (*ΔmdfA*; open inverted triangles), or cells deleted for *mdfA* and harboring *mdfA* integrated at the *amyE* locus (*ΔmdfA amyE*::*mdfA*; open triangles) (P*_spoIIQ_-lacZ* strains AHB1841, CFB530, and CFB541, respectively; P*_sspB_-lacZ* strains AHB324, SMB179, and SMB334, respectively). Error bars indicate ± standard deviations based on three or more independent experiments.

At the outset of this study, we sought to identify genes that drive the forespore toward metabolic dormancy, a poorly understood yet hallmark feature of sporulation. We reasoned that “metabolic differentiation” genes are likely to be expressed early under the control of σ^F^, given that the forespore soon thereafter becomes dependent upon an intercellular channel comprised of the SpoIIIAA-AH and SpoIIQ proteins, through which the mother cell provides small molecule metabolites (e.g. nucleotides, amino acids) to the forespore (Camp and Losick 2009; Morlot and Rodrigues 2018; Riley et al. 2021; Zeytuni and Strynadka 2020). The most recent of these studies (Riley et al. 2021), which was the first to directly demonstrate transport of small molecule metabolites via the SpoIIIAA-AH•SpoIIQ channel, also found that metabolic enzymes disappear from the forespore soon after asymmetric division, further supporting the notion that metabolic dormancy initiates early in forespore development (**Fig 1A**). Here we report the identification of the σ^F^-dependent *yjbA* gene (henceforth *mdfA* for *<u>m</u>*etabolic *<u>d</u>*ifferentiation *f*actor *<u>A</u>*) as a candidate factor involved in the forespore transition to metabolic dormancy. We demonstrate through genetic, biochemical, and structural approaches that the MdfA protein functions as a ClpCP adaptor protein. Our findings and those of an accompanying manuscript (Riley et al. 2024) together support a model in which MdfA induces ClpCP to degrade substrate proteins including metabolic enzymes in the forespore, the removal of which helps set the developing spore on a path to metabolic dormancy.

## RESULTS

### Deletion of the *mdfA* (*yjbA*) gene significantly stimulates forespore gene expression

To identify genes that help drive the forespore toward metabolic dormancy, we screened knockouts of 59 known or predicted σ^F^ target genes for phenotypes consistent with elevated forespore metabolic activity (**Fig S1, Table S5**). Most notably, deletion of the gene *yjbA*, which we have renamed here as *mdfA* (*m*etabolic *d*ifferentiation *f*actor *A*), stimulated both σ^F^- and σ^G^-dependent gene expression in the forespore. Cells deleted for *mdfA* displayed a ∼10-fold increase in the σ^F^-dependent expression of P*_spoIIQ_-lacZ* (**Fig 1B**), and a small but reproducible ∼1.6-fold increase in the σ^G^-dependent expression of *P_sspB_-lacZ* (**Fig 1C**), effects that were fully complemented by a wild-type copy of *mdfA*. Other σ^F^- and σ^G^-dependent reporters gave similar results (**Fig S2A, S2B**), indicating that forespore gene expression is broadly elevated in *ΔmdfA* cells. In contrast, σ^E^- and σ^K^-dependent mother cell gene expression was mostly unaffected (**Fig S2C, S2D**).

The *mdfA* gene has until now remained uncharacterized other than its assignment to the σ^F^ regulon by transcriptomic studies (Arrieta-Ortiz et al. 2015; Steil et al. 2005; Wang et al. 2006). We confirmed that *mdfA* is a bona fide σ^F^-target gene: *mdfA* upstream regulatory sequences, which include a putative σ^F^ consensus promoter sequence, activated reporter gene expression in the forespore during sporulation in a σ^F^-dependent manner (**Fig S3A,B,D**). The encoded MdfA protein is well-conserved within the endospore-forming *Bacilliaceae* family (**Fig S4**), but is absent from other bacteria including endospore-formers of the more distantly-related *Clostridia* genus. Homology searches yielded no evidence of more distantly related proteins or protein domains. A functional GFP-MdfA fusion protein expressed under native *mdfA* regulatory sequences localized to forespores and sometimes appeared enriched at the forespore periphery (**Fig S3C,E**). Interestingly, GFP-MdfA fluorescence was significantly dimmer than GFP alone expressed from the same *mdfA* regulatory sequences (**Fig S3D** vs. **Fig S3E**), suggesting that the MdfA protein may be targeted for degradation in the forespore.

Finally, we found that *ΔmdfA* mutants were modestly deficient for spore formation, producing ∼20-30% fewer heat-resistant spores than wild type cells in single-round sporulation assays (**Fig S5A**). The *ΔmdfA* mutant also displayed a competitive disadvantage relative to wild type cells during multiple rounds of growth and sporulation in co-culture (**Fig S5B, S5C**). This indicates that *mdfA* contributes to maximal sporulation efficiency, consistent with the sporulation-specific expression of *mdfA*, its effect upon forespore gene expression, and its conservation in the spore-forming *Bacilliaceae* family.

### Vegetative expression of *mdfA* causes cell filamentation and lysis by a mechanism requiring the ClpCP AAA+ protease

We next constructed a strain in which *mdfA* expression could be induced during vegetative growth. When grown on solid media with high concentrations of inducer, this strain formed small colonies that eventually lysed; at lower concentrations, growth appeared mostly normal (**Fig 2A**, **Fig S6A**). However, when observed by phase microscopy, cells from colonies grown at lower inducer concentrations were elongated, and individual lysed cells were also observed (**Fig 2B, Fig S6B**). Fluorescence microscopy revealed that the elongated cells were aseptate filaments that sometimes harbored evenly spaced nucleoids but were unable to form the FtsZ ring (“Z-ring”) responsible for the initiation of cell division in nearly all bacteria (**Fig S6C**) (Haeusser and Margolin 2016), likely accounting for the observed filamentous phenotype.

**Figure 2.**
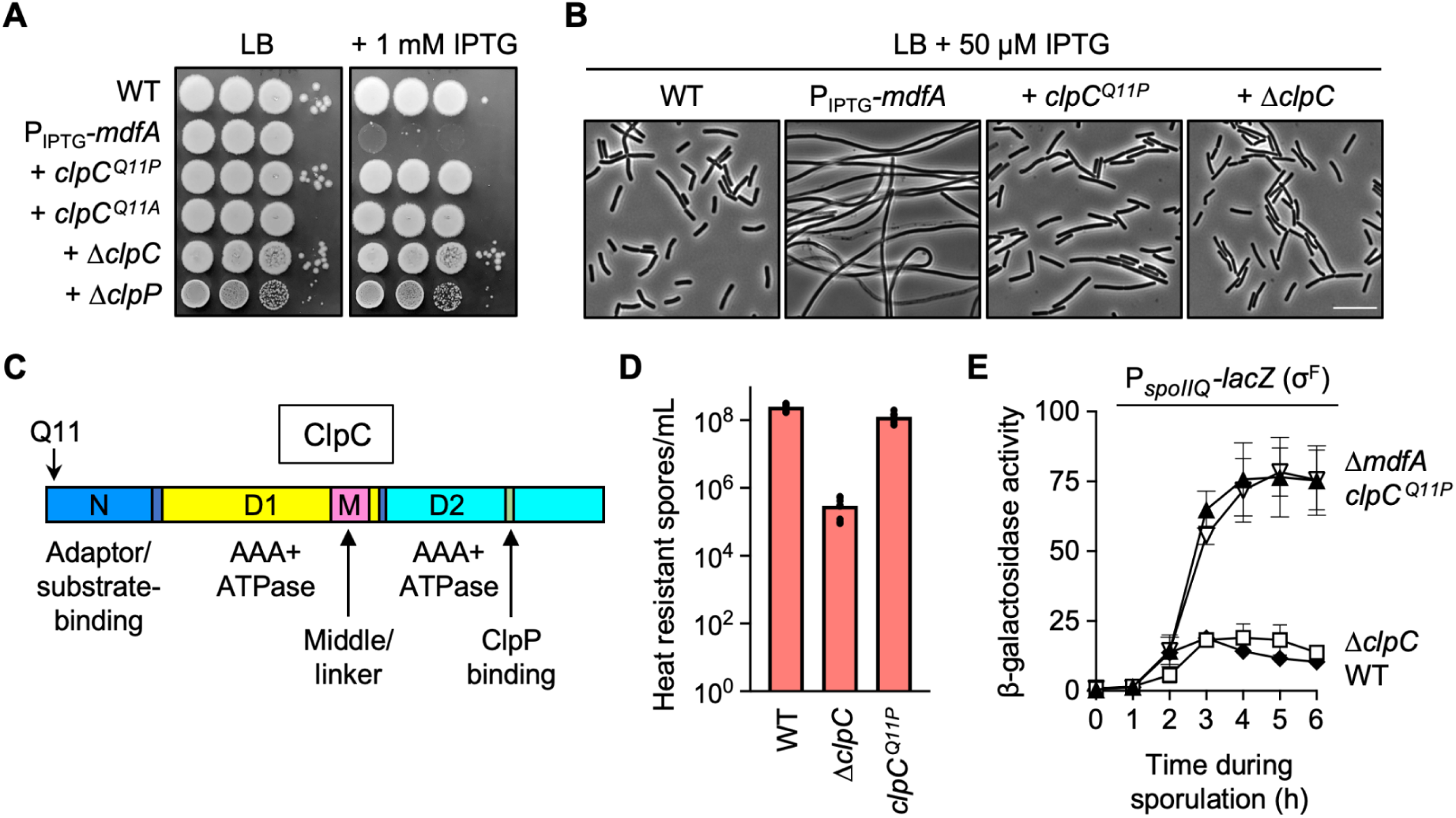
The *clpC^Q11P^* allele suppresses vegetative *mdfA* toxicity and, like *ΔmdfA*, stimulates σ^F^ activity during sporulation. Expression of *mdfA* during vegetative growth causes cell death at high inducer concentrations, and this toxicity is suppressed by mutations in *clpC* and *clpP*. Wild type cells and cells harboring an engineered IPTG-inducible *mdfA* gene construct (WT and P_IPTG_-*mdfA*; strains PY79 and CFB189, respectively), as well as P_IPTG_-*mdfA* cells harboring *clpC^Q11P^*, *clpC^Q11A^*, *ΔclpC*, or *ΔclpP* (strains SFB32, SFB38, CFB241, and CFB274, respectively), were grown to mid-logarithmic phase (OD_600_∼0.5) in LB media, adjusted to OD_600_=1, and 10-fold serial dilutions (10^-1^-10^-4^, left to right) were spotted onto LB plates with or without 1 mM IPTG. Plates were imaged after 24 h of growth at 37°C. Expression of *mdfA* during vegetative growth causes cell filamentation at lower inducer concentrations, and this filamentation is suppressed by *clpC^Q11P^*and *ΔclpC*. WT, P_IPTG_-*mdfA*, P_IPTG_-*mdfA clpC^Q11P^*, and P_IPTG_-*mdfA ΔclpC* cells (strains PY79, CFB189, SMB361, and CFB241, respectively) were grown for ∼24 h at 37°C on LB agar plates with 50 µM IPTG. Colonies were picked, diluted, and visualized by phase microscopy. Scale bar, 10 µm. Domain structure of the ClpC protein. Shown are the adaptor/substrate-binding N-domain (blue), AAA+ ATPase D1-domain (yellow), M-domain (magenta), and AAA+ ATPase D2-domain (cyan). The ClpP binding site within the D2-domain is also indicated in green. The location of the Q11 residue in the N-domain is labeled. Cells harboring the *clpC^Q11P^* allele, unlike those harboring *ΔclpC*, do not have a severe sporulation defect. WT, *ΔclpC*, and *clpC^Q11P^* cells (strains PY79, CFB270 and CFB282, respectively) were induced to sporulate for 24 h and the number of heat resistant spores per mL were determined. Individual data points are shown (n=6) and bars indicate the mean transformation efficiency for each strain. The *clpC^Q11P^* allele, like *ΔmdfA* but unlike *ΔclpC*, significantly stimulates σ^F^ activity during sporulation. β-Galactosidase production from the σ^F^-dependent P*_spoIIQ_-lacZ* reporter was monitored during sporulation of otherwise wild type cells (WT; closed diamonds), cells deleted for *mdfA* (*ΔmdfA*; open inverted triangles), cells deleted for *clpC* (*ΔclpC*; open squares) or cells harboring *clpC^Q11P^* (*clpC^Q11P^*; solid triangles) (strains AHB881, SMB189, CFB302, and CFB307, respectively). Error bars indicate ± standard deviations based on four replicates.

To identify the MdfA target(s) that drive cell filamentation and toxicity, we isolated extragenic suppressor mutations that restored normal growth to cells overexpressing *mdfA*. This genetic strategy was facilitated by the spontaneous appearance of such suppressors (**Fig 2A**, **Fig S6A, inset**), although we did modify our *mdfA* overproducing strain to minimize isolation of intragenic mutations in the expression construct itself (see **Supplemental Materials and Methods**). We found that extragenic suppression of *mdfA* vegetative toxicity was most commonly caused by mutations in the *clpC* gene. ClpC is a member of the Hsp100/Clp family of AAA+ chaperones that partners with the ClpP protease to degrade target proteins (see Introduction) (Sauer and Baker 2011). Notably, a strain deleted for *clpC* was protected against the colony lysis and cell filamentation phenotypes associated with *mdfA* expression (**Fig 2B-C**). Cells deleted for *clpP*, despite displaying a small colony phenotype even in the absence of inducer, were also immune to the toxic effects of *mdfA* expression (**Fig 2B**). These genetic data indicate that MdfA induces filamentation and toxicity in vegetative cells by a mechanism that requires the ClpCP proteolytic complex.

### The *mdfA*-specific *clpC^Q11P^* allele phenocopies *ΔmdfA* during sporulation

Most of our isolated *clpC* suppressor alleles harbored mutations predicted to severely compromise ClpC function (e.g. truncations, insertions) (**Table S1**). However, one isolated allele, *clpC^Q11P^*, fully suppressed *mdfA*-associated toxicity and filamentation through a single point mutation in the codon for Gln11 (Q11), switching it to proline (Q11P) (**Fig 2A-B**). Q11 is located in the ClpC N-domain responsible for binding substrates and/or adaptor proteins that recruit substrates to the protease complex (**Fig 2C**). Interestingly, unlike *ΔclpC* cells and more similar to *ΔmdfA* cells, *clpC^Q11P^* cells did not display a pleiotropic defect in sporulation efficiency (**Fig 2D**) (Meeske et al. 2016), suggesting that *clpC^Q11P^* is likely to be a *mdfA*-specific *clpC* allele. Consistent with a specific interaction at Q11, we found that its substitution for alanine (Q11A) also suppressed *mdfA*-associated toxicity (**Fig 2A**).

We reasoned that we could use the *mdfA*-specific *clpC^Q11P^* allele to assess whether MdfA mediates its effects on forespore physiology in collaboration with ClpCP, which is known to be present and active in the forespore during sporulation (Kain et al. 2008), without the complication of other pleiotropic defects of a *ΔclpC* mutant. Strikingly, *clpC^Q11P^* caused the same stimulation of forespore gene expression as observed in *ΔmdfA* cells (**Fig 2E**). In contrast, the *ΔclpC* mutant did not exhibit any stimulation of σ^F^ activity, but rather a modest delay (**Fig 2E**), consistent with known pleiotropic sporulation defects of *ΔclpC* cells, including delayed entry (Meeske et al. 2016). These findings further support the notion that *clpC^Q11P^*is a *mdfA*-specific *clpC* allele. Finally, no additional σ^F^ activity was observed in a Δ*mdfA clpC^Q11P^* double mutant (data not shown), suggesting that *mdfA* and *clpC* likely operate in the same pathway to regulate forespore gene expression.

### MdfA interacts with the ClpC N-domain in a Q11-dependent manner

To determine whether MdfA directly interacts with the ClpC N-domain (ClpC^N^), we purified recombinant versions of each protein and analyzed their association through several approaches. When MdfA and ClpC^N^ were combined at different molar ratios and subjected to analytical size exclusion chromatography (SEC), formation of a higher molecular weight MdfA-ClpC^N^ complex was evident (**Fig S7**). Next, we used nuclear magnetic resonance (NMR) to collect heteronuclear single quantum coherence (HSQC) spectra of ^15^N-labeled ClpC^N^ or ^15^N-labeled ClpC^N,Q11P^ alone and in combination with increasing molar ratios of unlabeled MdfA (**Fig S8**). The addition of MdfA altered the ClpC^N^ spectrum, indicating complex formation; in contrast, the ClpC^N,Q11P^ spectrum remained stable in the presence of MdfA at four-fold molar excess, suggesting that the Q11P substitution disrupted interaction (**Fig S8C** vs. **Fig S8D**).

Finally, we employed isothermal titration calorimetry (ITC) to confirm a weak binding interaction

between MdfA and ClpC^N^ (K_d_ = 3.79 µM), while the ClpC^N,Q11P^ variant was unable to interact (**Fig 3A**).

**Figure 3.**
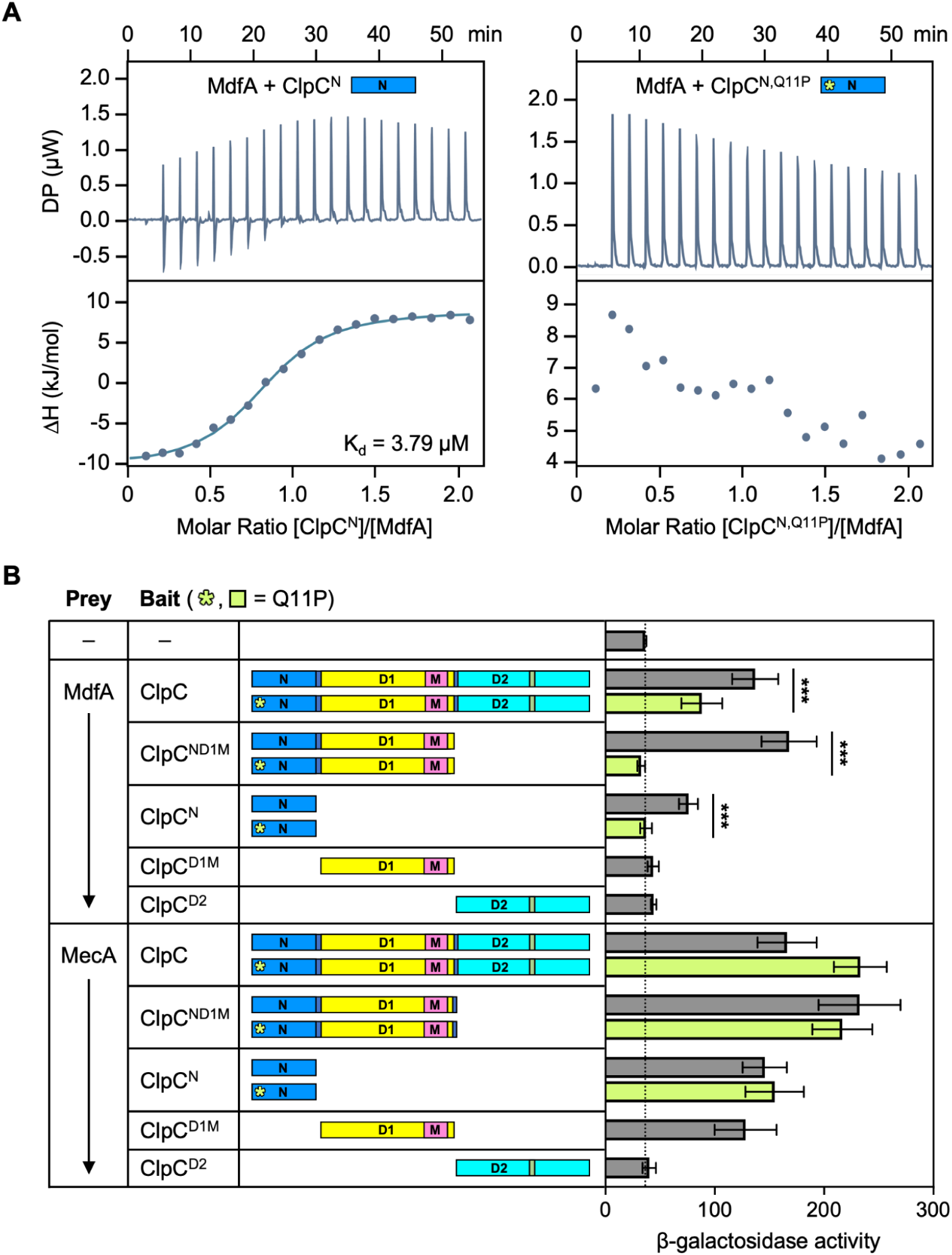
MdfA interacts with the ClpC N-domain in a Q11-dependent manner. Isothermal titration calorimetry (ITC) experiments indicate that MdfA and ClpC^N^ (left), but not MdfA and ClpC^N,Q11P^ (right) form a complex in vitro. Twenty injections of 2 μl MdfA at a concentration of 700 μM were added to ClpC constructs at 70 μM in the reaction cell. The heat of MdfA dilution into buffer was subtracted from the data. Raw data is shown in upper panels, binding isotherm in lower panels. Fitted data for MdfA-ClpC^N^ interaction: ΔH = −19.5 ± 0.6 kJ/mol; K_D_ = 3.79 ± 0.50 µM; N = 0.82 ± 0.01 sites. MdfA and ClpC interact in a transcription-based bacterial two-hybrid assay in a manner that depends upon the ClpC N-domain and Q11. In this assay, interaction of “Bait” and “Prey” proteins, fused respectively to the bacteriophage λ cI protein (λcI) and the N-terminal domain of the α subunit of RNA polymerase (α), drives expression of a *lacZ* reporter gene under the control of a test promoter in *E. coli*, leading to production of β-galactosidase (Dove and Hochschild 2004). ClpC or variants thereof were expressed from plasmids as fusions to λcI (“Bait”), while MdfA or the well-characterized ClpC adaptor protein MecA were expressed as fusions to α (“Prey”). The full list of bacterial two-hybrid bait and prey plasmids is provided in **Supplemental Table S2**. Variants of ClpC (ClpC^ND1M^, ClpC^N^, ClpC^D1M^, and ClpC^D2^) harboring one or more of its domains and/or the Q11P substitution are depicted in a manner derived from **Fig 2C**. The Q11P substitution is indicated by a pale green asterisk, and data collected from pairs including a ClpC Q11P variant are shaded pale green. The dotted line represents background β-galactosidase activity in a negative control strain expressing λCI and the α subunit of RNA polymerase alone (depicted as “–” for both prey and bait). Data are plotted as mean values, with error bars indicating ± standard deviation (n≥5 replicate values). \*\*\**p* < 0.0001, unpaired *t* test with Welch’s correction.

To further explore the interaction between ClpC and MdfA, we made use of a transcription activation-based bacterial two-hybrid assay (Dove and Hochschild 2004). When full-length ClpC and MdfA were paired as bait and prey, we observed a nearly 4-fold stimulation of reporter gene expression relative to a negative control strain (**Fig 3B**), indicating an interaction. MdfA also interacted with ClpC variants harboring the N-domain (ClpC^N^ and ClpC^ND1M^), but not those lacking the N-domain (ClpC^D1M^ and ClpC^D2^) (**Fig 3B**), indicating that the ClpC N-domain is both necessary and sufficient for interaction with MdfA in this assay. However, we noted that MdfA interacted more robustly with ClpC^ND1M^ relative to ClpC^N^ (4-fold vs. 2-fold stimulation of reporter gene expression, respectively), suggesting that MdfA may also make stabilizing contacts with the ClpC D1- and/or M-domains (**Fig 3B**). The interaction of MdfA with all ClpC variants harboring the N-domain was significantly reduced (full-length ClpC) or became undetectable (ClpC^ND1M^ and ClpC^N^) when the Q11P substitution was introduced (**Fig 3B**), indicating that ClpC Q11 is required for (robust) interaction with MdfA.

Finally, we used the bacterial two-hybrid assay to compare the interaction of ClpC with MdfA to the interaction of ClpC with its well-characterized adaptor protein MecA (Schlothauer et al. 2003; Wang et al. 2011). As predicted from previous studies (Kirstein et al. 2006; Wang et al. 2011), MecA interacted with ClpC variants harboring the N- or D1M-domains (ClpC, ClpC^N^, ClpC^ND1M^, and ClpC^D1M^) (**Fig 3B**). The Q11P substitution had no effect on the MecA-ClpC interaction (**Fig 3B**), revealing that the ClpC N-domain residue Q11 is specifically required for interaction with MdfA.

### MdfA induces oligomerization and ATPase activity of full-length ClpC in vitro

The genetic and physical interaction between MdfA and ClpC suggested to us a working model in which MdfA functions as an adaptor protein, binding to and modulating the activity of the ClpCP proteolytic complex in the forespore. To test this, we first implemented SEC to determine whether MdfA could induce the formation of higher oligomeric complexes of ClpC in vitro. We used a ClpC E280A/E618A double walker B variant (ClpC^DWB^), which binds but does not hydrolyze ATP and is known to form more stable higher oligomeric complexes with its adaptor protein MecA (Kirstein et al. 2006). Strikingly, ClpC^DWB^ formed large oligomeric complexes when mixed with MdfA in equimolar concentrations in the presence of ATP, although we did note that the complexes were smaller than those induced by MecA (**Fig 4A-B**). Given that MdfA is larger than MecA (29 kDa vs 26 kDa, respectively), this may suggest that the observed complex is not a 6:6 stoichiometry of ClpC:MdfA as is the case for ClpC:MecA (Wang et al. 2011).

**Figure 4.**
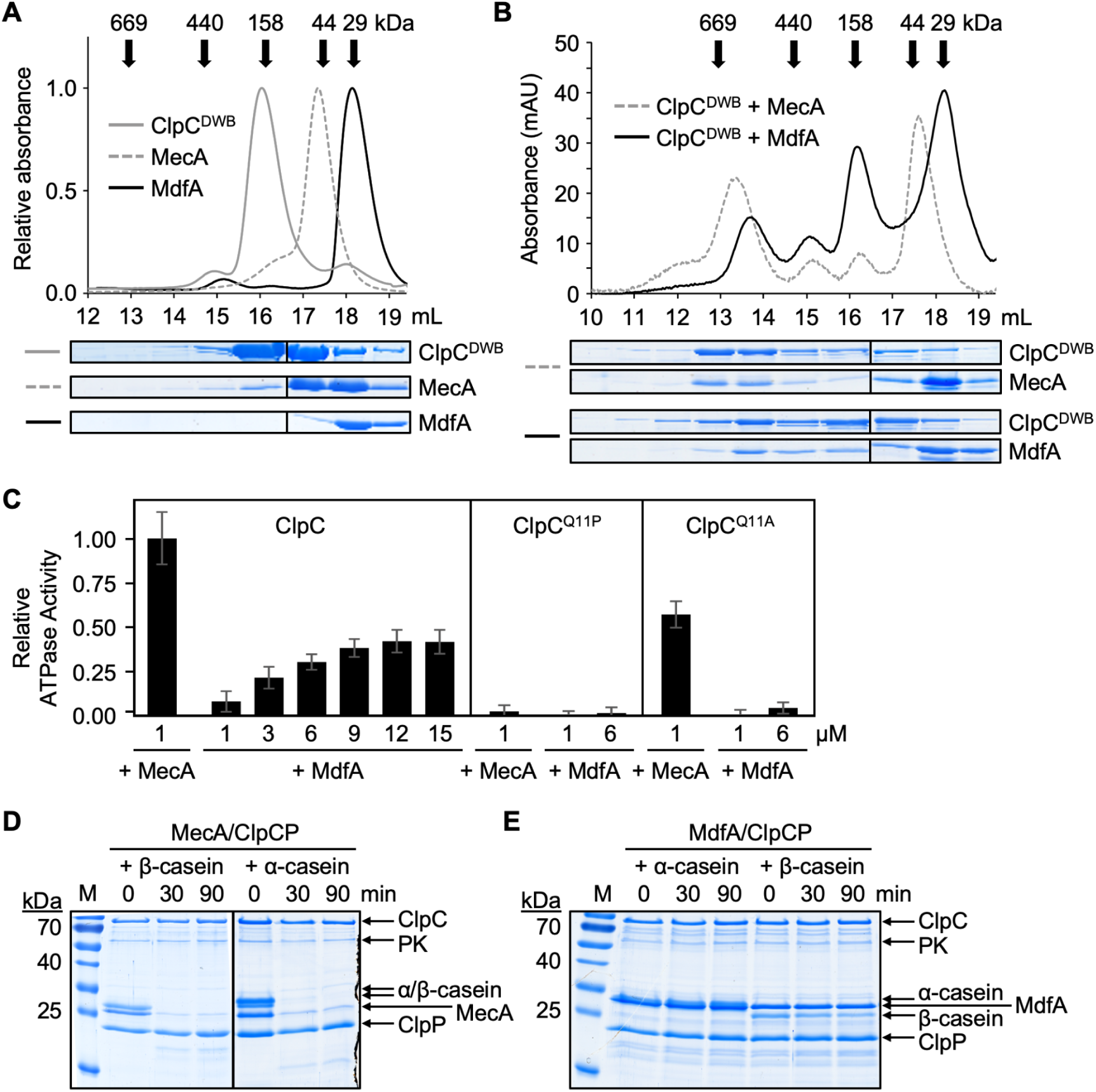
MdfA induces oligomerization and ATPase activity of full-length ClpC in vitro, but does not direct degradation of a test substrate. (A,B) Like MecA, MdfA induces the formation of stable higher oligomeric complexes with a ClpC double Walker B variant (ClpC^DWB^) in the presence of ATP, as detected by size exclusion chromatography. The elution profiles of all proteins **(A)** alone and **(B)** in combinations (as indicated; 10 µM each) were detected by absorbance at 280 nm. The positions and molecular weights of native marker proteins are indicated with black arrows. Elution fractions were concentrated and analyzed by SDS-PAGE followed by Coomassie staining, the results of which are shown below each relevant fraction. Like MecA, MdfA induces the ATPase activity of ClpC. The ATPase activity of ClpC, ClpC^Q11P^, or ClpC^Q11A^ (1 µM each) was determined in the presence of MecA (1 µM) or MdfA (1-15 µM, as indicated). All ATPase activities were normalized to that of ClpC induced by MecA. Error bars indicate ± standard deviations based on at least three replicates. **(D,E)** Unlike MecA, MdfA does not induce ClpCP-mediated degradation of itself nor α/β-casein. Samples from in vitro degradation reactions harboring the indicated proteins (1 µM each) as well as an ATP regeneration system (pyruvate kinase [PK]) were taken at 0, 30, and 90 min and analyzed by SDS-PAGE followed by Coomassie staining.

A second hallmark of ClpC adaptor proteins, including MecA, is their ability to induce ClpC ATPase activity (Schlothauer et al. 2003). We found that MdfA induced the ATPase activity of ClpC, albeit to a lesser extent than MecA (**Fig 4C**). Increasing the relative concentration of MdfA led to a maximum ClpC ATPase rate that was approximately one third of that induced by MecA. ClpC^Q11P^ ATPase activity could not be induced by either MecA or MdfA, suggesting that this mutant is nonfunctional under these in vitro conditions. However, a ClpC^Q11A^ variant was induced by MecA but not by MdfA. (Note that the *clpC^Q11A^*mutant is indistinguishable from the *clpC^Q11P^* mutant in suppressing *mdfA*-induced toxicity during vegetative growth [**Fig 2A**].) Finally, given that MdfA forms a higher oligomeric complex with ClpC and induces its ATPase activity, we wondered whether ClpC in complex with its cognate protease, ClpP, would degrade MdfA and/or a test substrate in vitro. Unlike MecA, however, MdfA was not degraded by ClpCP nor did it stimulate α/β-casein degradation (**Fig 4D-E**) (Schlothauer et al. 2003).

### Co-crystal structure of ClpC^N^ and MdfA

Finally, we solved the X-ray crystal structure of MdfA in complex with ClpC^N^, a 1:1 heterodimer, at 2.0 Å resolution (deposited as PDB: 8B3S; **Fig 5A**). ClpC^N^ overlays with an RMSD of 1.56 Å to a previously reported ClpC^N^ structure (PDB: 2Y1Q) (Wang et al. 2011). MdfA consists of an N-terminal domain (MdfA^N^, residues 1-135) and a C-terminal domain (MdfA^C^, residues 143-250), connected by a disordered linker. MdfA^N^ includes a small antiparallel beta-barrel and an isolated beta hairpin surrounded by three alpha-helices, while MdfA^C^ is an alpha-helical bundle. Both MdfA domains make contacts with ClpC^N^, fixing their positions relative to one another. The structure of MdfA appears to be novel, as we were unable to find significant structural homologs for the full-length protein. When the MdfA domains were analyzed individually, the N-domain still returned no significant structural homologs, whereas the C-domain exhibited structural similarity to a variety of helical-bundle-containing proteins including members of the eukaryotic death domain (DD) superfamily, such as RIP, CARD and Pyrin type proteins (Park et al. 2016).

**Figure 5.**
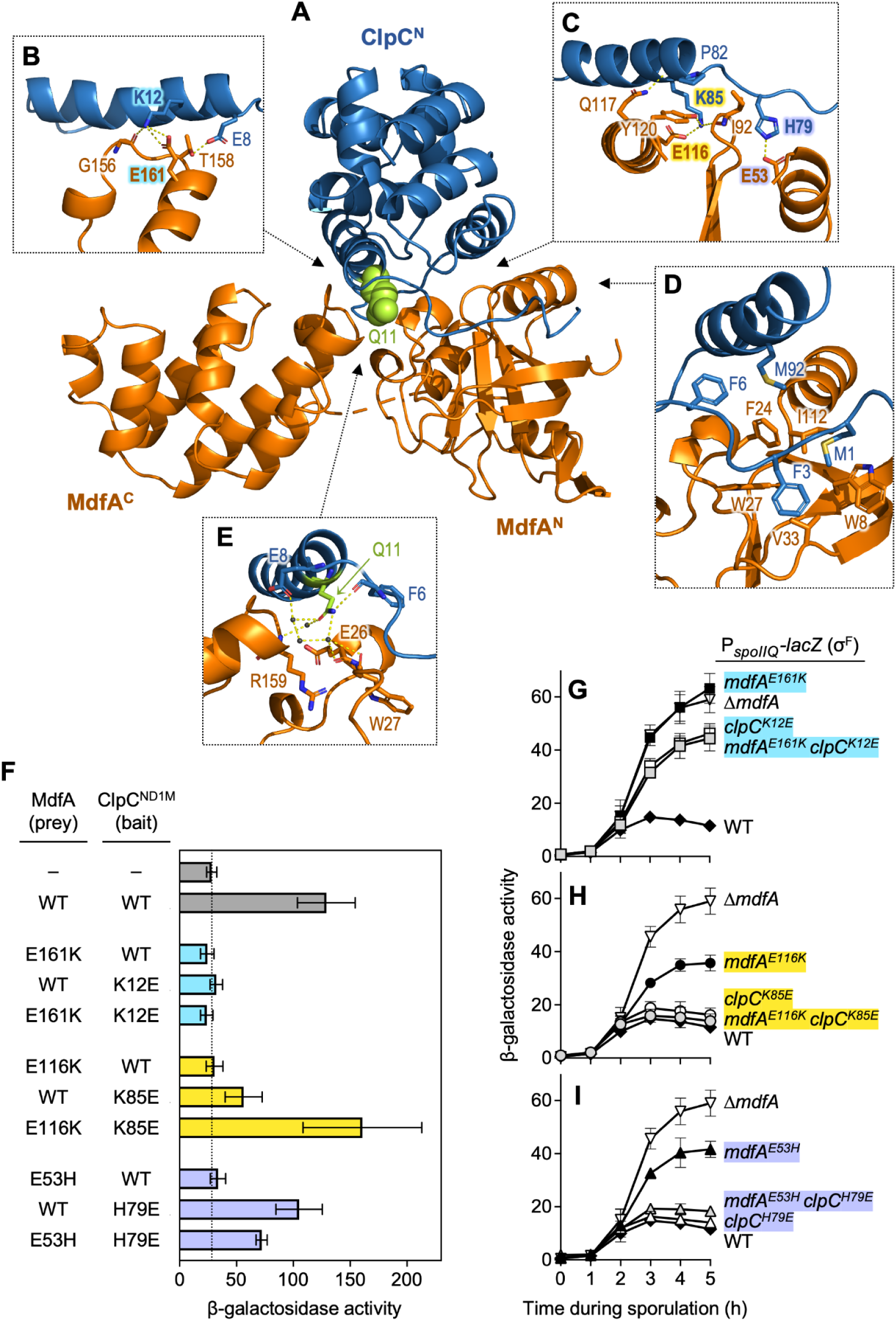
A co-crystal structure of MdfA and ClpC^N^. (A) Complete X-ray crystal structure of MdfA (orange) in complex with ClpC^N^ (blue) with Q11 highlighted (lime green space fill) solved to 2.0 Å resolution. **(B-E)** Zoomed views of the interface between MdfA (orange) and ClpC^N^ (blue). Three pairs of charged residues predicted to form electrostatic interactions between the two proteins are highlighted: ClpC K12 and MdfA E161, ClpC K85 and MdfA E116, and ClpC H79 and MdfA E53. **(F)** Mutational analysis of the MdfA-E161/ClpC-K12, MdfA-E116/ClpC-K85, and MdfA-E53/ClpC-H79 electrostatic interactions in MdfA-ClpC^ND1M^ binding in the bacterial two-hybrid assay. See Figure 3 legend for more details. The full list of bacterial two-hybrid bait and prey plasmids is provided in **Supplemental Table S2**. The dotted line represents background β-galactosidase activity in a negative control strain (depicted as “–” for both prey and bait). Data are plotted as mean values, with error bars indicating ± standard deviation (n≥4 replicate values). **(G-I)** Mutational analysis of the **(G)** MdfA-E161/ClpC-K12, **(H)** MdfA-E116/ClpC-K85, and **(I)** MdfA-E53/ClpC-H79 electrostatic interactions during sporulation. β-Galactosidase production from the σ^F^-dependent P*_spoIIQ_-lacZ* reporter was monitored during sporulation of otherwise wild type cells (WT; closed diamonds), cells deleted for *mdfA* (*ΔmdfA*; open inverted triangles), or cells harboring the indicated mutations in *mdfA* and/or *clpC* (closed squares/circles/triangles indicate *mdfA* single mutants, open squares/circles/triangles indicate *clpC* single mutants, and shaded squares/circles/triangles indicate *mdfA clpC* double mutants). Strains were as follows: WT (AHB881), *ΔmdfA* (SMB189), *mdfA^E161K^* (SYB10), *clpC^K12E^* (RFB32), *mdfA^E161K^ clpC^K12E^*(AHB6234), *mdfA^E116K^* (AHB6220), *clpC^K85E^* (AHB6211), *mdfA^E116K^ clpC^K85E^* (AHB6229), *mdfA^E53H^* (AVB6), *clpC^H79E^* (AHB6207), *mdfA^E53H^ clpC^H79E^* (AHB6231). Error bars indicate ± standard deviations based on three or more replicates. The same WT and *ΔmdfA* data is shown in all three panels for clarity.

The interface between MdfA and ClpC^N^ covers 1360 Å^2^, and involves a combination of hydrophobic (**Fig 5D**) and electrostatic (**Fig 5B,C,E**) interactions, the latter including three acidic side chains on MdfA (E161, E116, and E53) that are within bonding distance of three corresponding basic side chains on ClpC (K12, K85, and H79, respectively). MdfA and ClpC mutant proteins with switched charges at these positions displayed reduced interaction with their wild type partner proteins in the bacterial two-hybrid assay (**Fig 5F**). We speculate that residual interaction of ClpC^K85E^ and ClpC^H79E^ with MdfA could be due to compensatory interactions with nearby charged and polar side chains in ClpC. For two of the three electrostatic pairs, interaction was fully (for MdfA-E116/ClpC-K85) or partially (for MdfA-E53/ClpC-H79) restored if the two side chains were simultaneously charge swapped (**Fig 5F**), highly suggestive of a direct interaction. When these mutations were introduced into the native *mdfA* and *clpC* genes in *B. subtilis*, MdfA/ClpC function during sporulation was disrupted as evidenced by stimulated σ^F^ activity (**Fig 5G-I**). The two exceptions were *clpC^K85E^* and *clpC^H79E^*, which appeared fully functional, likely reflecting the residual ability of their encoded proteins to interact with MdfA (**Fig 5F**). Still, these mutants restored wild type function to the *mdfA^E116K^* and *mdfA^E53H^* mutants, respectively (**Fig 5H and Fig 5I**), consistent with the restored interaction observed for these double charge swaps (**Fig 5F**). We noted that the *clpC^K12E^* mutant was likewise able to rescue some functionality to the *mdfA^E161K^* mutant during sporulation (**Fig 5G**), despite no evidence of a restored interaction (**Fig 5F**), hinting at some contact between these side chains in vivo. Finally we tested the effect of these mutations on *mdfA* toxicity during vegetative growth, the results of which are consistent with those described above (**Fig S9**). In this assay, the *mdfA^E161K^ clpC^K12E^* double mutant showed robust restoration of toxicity relative to the single mutants, lending stronger support to an interaction between MdfA-E161 and ClpC-K12 in vivo.

Interestingly, ClpC Q11 does not bind directly to MdfA in our co-crystal structure, but rather does so indirectly via a network of water molecules (**Fig 5E**); as noted above, the adjacent residue K12 appears to directly interact with MdfA (**Fig 5B**). When queried, AlphaFold Multimer (Evans et al. 2022) was unable to predict the same binding interface, and instead positioned ClpC^N^ with low confidence on the opposite surface of MdfA. The ClpC^N^ binding surface occupied by MdfA in our co-crystal structure is distinct from that bound by other known ClpC^N^ protein interactors (MecA and phosphoarginine-modified proteins) (Wang et al. 2011; Trentini et al. 2016), but is analogous to the surface of *M. tuberculosis* ClpC1 that is bound by various antibiotics (cyclomarin A, ecumicin, rufomycin) (Vasudevan et al. 2013; Wolf et al. 2020, 2019) (**Fig 6**).

**Figure 6.**
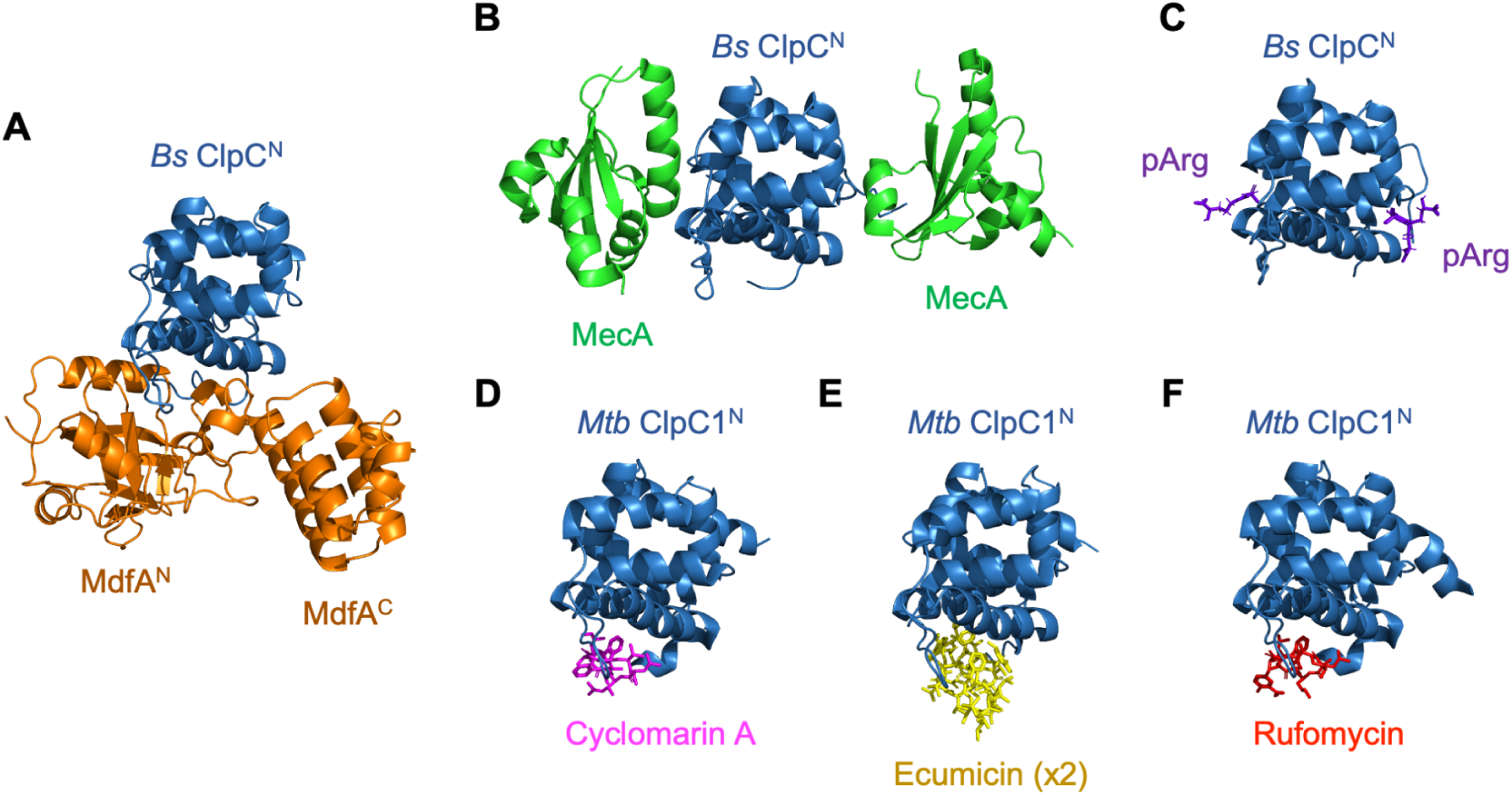
MdfA binds ClpC^N^ in a location distinct from that bound by the *B. subtilis* ClpC adaptor MecA and phosphoarginine (pArg) peptides, but similar to the surface of *M. tuberculosis* ClpC1^N^ bound by the antibiotics Cyclomarin A, Ecumicin, and Rufomycin. (A-C) Co-crystal structures of the *B. subtilis* ClpC N-domain bound to **(A)** MdfA (PDB: 8B3S; this study), **(B)** MecA (PDB: 3PXG) (Wang et al. 2011), and **(C)** phosphoarginine (pArg; PDB: 5HBN) (Trentini et al. 2016). **(D-F)** Co-crystal structures of the *M. tuberculosis* ClpC N-domain bound to **(D)** cyclomarin A (PDB: 3WDC) (Vasudevan et al. 2013), **(E)** ecumicin (PDB: 6PBS) (Wolf et al. 2020), and **(F)** rufomycin (PDB: 6CN8) (Wolf et al. 2019).

To examine the effect of the *clpC^Q11P^* mutation, we also solved the crystal structure of ClpC^N,Q11P^ to 1.2 Å resolution (deposited as PDB: 8OTK). It closely resembles previous ClpC structures with the exception of a slight kink in the first alpha-helix at the location of the Q11P substitution (**Fig S9**).

Despite extensive efforts we were unable to crystallize MdfA in isolation, which we believe is a consequence of the mobility conferred on MdfA by the flexible linker that connects its two domains. AlphaFold (Jumper et al. 2021) predicts the MdfA monomer as being essentially structurally identical to how it appears in the complex crystal structure (overlaying with RMSD = 1.0), including the orientation of the two domains relative to one another. However, when we performed small-angle X-ray scattering (SAXS) on MdfA, the results were incompatible with the rigid AlphaFold and complex crystal structures (**Fig S10**), supporting the idea that the N- and C-domains of MdfA can move relative to one another via the linker in the absence of a binding partner.

## DISCUSSION

Here we report how a search for factors involved in metabolic differentiation of the developing *B. subtilis* spore led us to discover that the forespore-expressed protein MdfA (formerly YjbA) functions as a ClpCP adaptor protein. Through genetic, biochemical, and biophysical approaches, we demonstrate that MdfA directly interacts with the ClpC N-domain and induces ClpC oligomerization and ATPase activity. We further solved a co-crystal structure of MdfA bound to the ClpC N-domain, revealing that MdfA binds to a surface of the ClpC N-domain that is unique among its known adaptor proteins but that overlaps with the homologous binding sites of antibiotics known to misregulate the *M. tuberculosis* ClpC1 protein (**Fig 6**) (Trentini et al. 2016; Vasudevan et al. 2013; Wang et al. 2011; Wolf et al. 2020, 2019).

An open question remaining from our study is the identity of the protein(s) targeted by MdfA-activated ClpCP in the forespore and how their degradation promotes the transition to metabolic dormancy. Recently, Riley et al. (2021) demonstrated that metabolic enzymes in the forespore become depleted soon after asymmetric division, which raises the intriguing idea that these metabolic enzymes are the substrates for MdfA-activated ClpCP. Now, in an accompanying study, Riley, Lyda and colleagues report their independent discovery that ClpC and MdfA are indeed required for the degradation of a subset of these metabolic enzymes, along with other proteins including transcriptional regulators (Riley et al. 2024). Satisfyingly, they show that amino acid substitutions in MdfA and ClpC that disrupt and then restore the interaction of these two proteins, as predicted by our co-crystal structure and confirmed in our study, disrupt and then restore degradation of the metabolic enzyme CitZ. Collectively the results of our two complementary studies support a model in which MdfA functions as a forespore-specific adaptor protein for ClpCP: triggering its assembly and activating it to degrade numerous proteins including metabolic enzymes, in turn setting the forespore on its path to metabolic dormancy.

### How does MdfA bind and activate ClpC?

The MdfA adaptor joins a growing list of proteins and small molecules that bind to and activate ClpC. On its own, ClpC forms an inactive resting decamer state (Carroni et al. 2017; Morreale et al. 2022; Taylor et al. 2022), but binding partners can drive it out of this resting state to an active hexamer or even tetramer of hexamers (Morreale et al. 2022; Maurer et al. 2019; Carroni et al. 2017). The best studied ClpC-binding protein is the MecA adaptor, which activates *B. subtilis* ClpC by promoting its assembly into a functional hexamer, with MecA binding in a 1:1 ratio with ClpC N-domains and M-domain linkers (Carroni et al. 2017; Kirstein et al. 2006; Wang et al. 2011). Substrate proteins that have been phosphorylated by McsB on arginine residues can also promote assembly of active ClpC hexamers (Trentini et al. 2016).

Here we have shown that MdfA triggers ClpC hexamerization and ATPase activity, analogous to the *B. subtilis* ClpC adaptor MecA (Wang et al. 2011). However, our solved ClpC^N^-MdfA structure is sterically incompatible with published structures of the ClpC hexamer formed through binding with MecA. As shown in **Figure 7**, when we dock MdfA onto the ClpC hexamer using the ClpC N-domain for alignment, we observe clashes between MdfA and the ClpC M-domain. However, the ClpC N-domain is separated from the rest of the protein by a 15-residue flexible linker, which likely allows its repositioning (Wang et al. 2011). The M-domain has also been demonstrated to move by 6.0 Å during substrate processing, which might also accommodate MdfA binding (Liu et al. 2013). It is interesting to consider the possibility that, like MecA, MdfA may interact with the M-domain, which could explain our bacterial two-hybrid data that MdfA binds better to ClpC^ND1M^ than to ClpC^N^ alone. This would also suggest a mechanism by which MdfA promotes hexamerization and activation of ClpC, given that assembly of the inactive ClpC state requires head-to-head M-domain interactions (Carroni et al. 2017). Future determination of the full MdfA-ClpC assembled structure, likely using cryo-electron microscopy to observe multiple states, will prove informative both in observing the structural differences as compared to other adaptors but also in understanding the mechanism of MdfA-induced ClpC activation.

**Figure 7.**
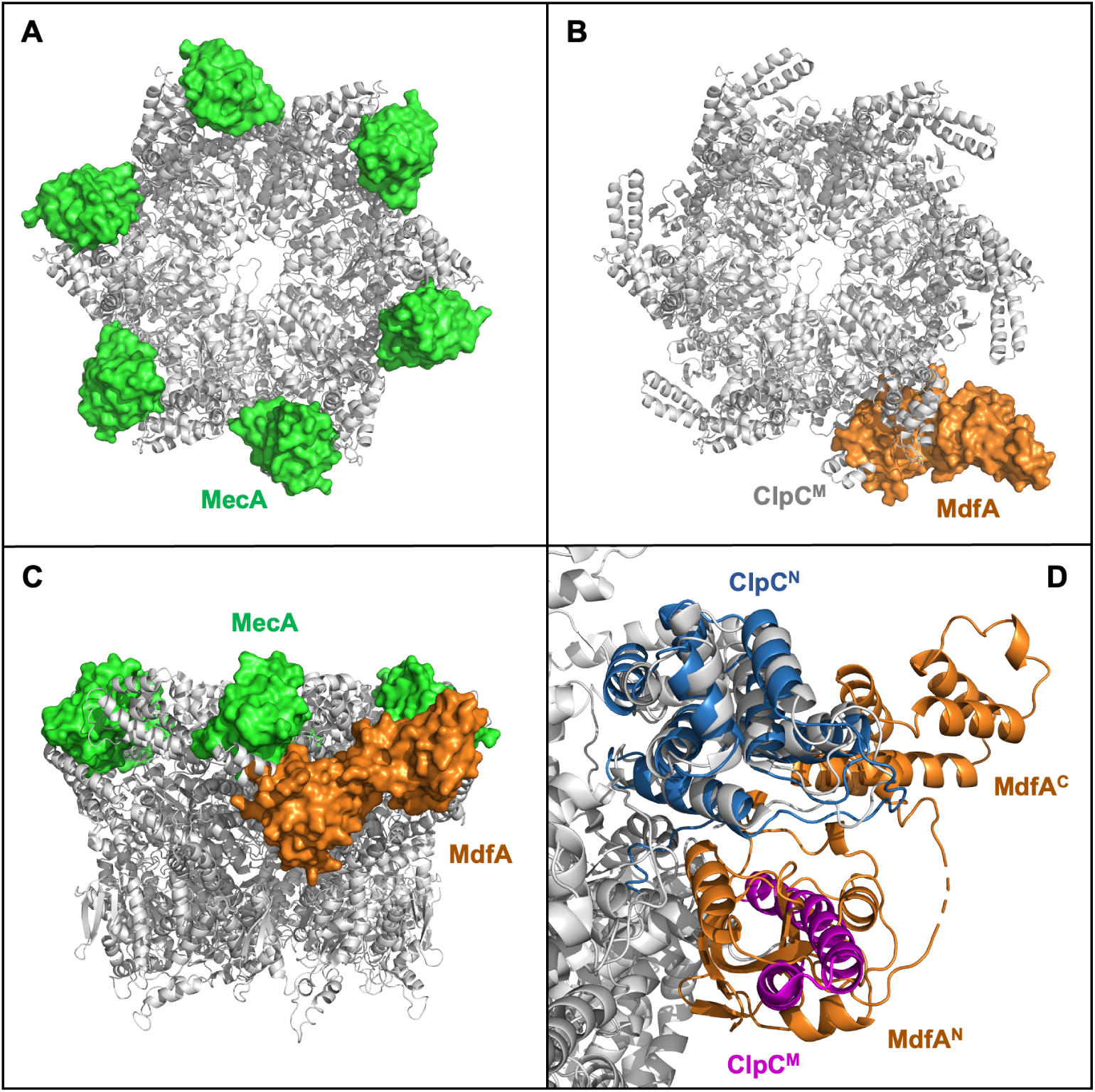
Modeling MdfA onto the ClpC hexamer indicates potential steric clash with the ClpC M-domain. (A) Co-crystal structure (PDB: 3J3T) of *B. subtilis* MecA (green) in a hexameric complex with ClpC (gray) (Liu et al. 2013). **(B,C,D)** Alternate views of the MdfA/ClpC^N^ crystal complex (PDB: 8B3S, this work) aligned with ClpC^N^ in the full length ClpC hexamer shown in (A), indicating that MdfA (orange) and MecA (green) occupy different binding positions on ClpC^N^ **(D)** Zoomed view showing the steric clash between MdfA (orange) and the ClpC M-domain (magenta). The overlay of ClpC^N^ from the hexamer (gray) and the MdfA/ClpC^N^ co-crystal structure (blue) is shown in (D) but removed from images (B) and (C) for clarity.

The significance of MdfA binding to a surface of ClpC^N^ that overlaps with the homologous binding sites of three antibiotics that target *M. tuberculosis* ClpC1 is unclear. Even among these antibiotics, binding at this surface results in different effects upon ClpC1 and the ClpC1-ClpP1-ClpP2 (ClpC1P1P2) proteolytic complex. The cyclic peptide cyclomarin A increases both the ATPase of ClpC1 and proteolytic activity of ClpC1P1P2, such that substrate specificity is relaxed (Maurer et al. 2019; Taylor et al. 2022); in contrast, the cyclic peptides ecumicin and rufomycin decrease ClpC1P1P2 proteolytic activity, while either increasing (ecumicin) or leaving unaffected (rufomycin) ClpC1 ATPase activity (Gao et al. 2015; Choules et al. 2019). Like cyclomarin A and ecumicin, we have found that MdfA stimulates ClpC ATPase activity, but have so far been unable to demonstrate the ability of MdfA to stimulate ClpCP proteolysis in vitro. We nevertheless infer that MdfA exerts its in vivo effects by promoting (as opposed to inhibiting) ClpCP-mediated proteolysis, given our genetic finding that deletion of either *clpC* or *clpP* suppresses vegetative toxicity of *mdfA* overexpression. It is tempting to speculate that MdfA, like cyclomarin A, might constitutively activate ClpCP-dependent proteolysis, but our finding that MdfA could not stimulate ClpCP to degrade casein in vitro may argue against this.

### How does MdfA impact forespore development?

We originally flagged *mdfA* as a gene that may help drive the forespore toward metabolic dormancy given that gene expression in the forespore (both σ^F^- and σ^G^-dependent) is significantly stimulated in its absence, a phenotype that we reasoned may reflect increased metabolic activity (e.g. increased availability of metabolic products such as nucleotides and amino acids). The stabilization of metabolic enzymes in *ΔmdfA* forespores reported in the accompanying study supports this explanation (Riley et al. 2024). However, they also found that the σ^F^ protein itself is stabilized in *ΔmdfA* forespores, providing a more direct explanation for increased levels of σ^F^-dependent gene expression. This could also explain increased σ^G^-dependent gene expression given that σ^F^ activates transcription of *sigG*, the gene encoding σ^G^ (Mearls et al. 2018; Sun et al. 1991). We speculate that forespore gene expression is stimulated in *ΔmdfA* cells due to a combination of these two mechanisms: stabilized σ^F^ and increased metabolic capacity.

Finally, we speculate that degradation of proteins by MdfA-activated ClpCP must not be the only mechanism by which the forespore progresses toward metabolic dormancy, given that we observed only a modest defect in heat resistant spore formation by cells lacking *mdfA*. An exciting challenge for future work will be to identify the redundant pathways that drive metabolic dormancy of the developing spore.

## MATERIALS AND METHODS

### General *B. subtilis* methods

All *B. subtilis* strains were isogenic with the laboratory strain PY79 (Youngman et al. 1984) and were typically propagated in lysogeny broth (LB), in liquid culture or on solid plates with 1.6% agar. Tables of strains, plasmids, primers, and synthetic DNA fragments used in this study, as well as details of strain and plasmid construction, are provided as **Supplemental Materials**. To quantify spore formation, cells were induced to sporulate by nutrient exhaustion for 24 h at 37°C in Difco sporulation medium (DSM) (Nicholson and Setlow 1990; Schaeffer et al. 1965), after which the number of colony forming units/mL that survived heat treatment (80°C for 20 min) was determined. For other experiments, including β-galactosidase activity assays and fluorescence microscopy, sporulation was induced by the resuspension method (Nicholson and Setlow 1990; Sterlini and Mandelstam 1969). Cells were collected at intervals by centrifugation and either processed immediately or stored at -80°C for later analysis.

β-galactosidase activity in samples from *B. subtilis* strains harboring *lacZ* reporters was measured as previously described (Camp and Losick 2009). Details regarding microscopy are provided in **Supplemental Materials and Methods**.

### Bacterial two-hybrid assay

Bacterial two-hybrid assays (Dove and Hochschild 2004) were carried out in the *E. coli* strain FW102 O_L_2-62 (Deaconescu et al. 2006). FW102 O_L_2-62 strains harboring the appropriate bait and prey plasmids (**Supplemental Table S2**) was grown overnight (∼16 h) at 37°C in 3 mL LB supplemented with the necessary antibiotics and 10 µM IPTG to induce expression of the bait and prey protein fusions. The next morning, 30 µL of the overnight culture was back-diluted into 3 mL fresh LB supplemented with the necessary antibiotics and 10 µM IPTG and grown at 37°C to OD_600_ ∼0.5, at which point β-Galactosidase activity was measured by a hybrid protocol based on Thibodeau et al. (2004) and Camp and Losick (2009). 100 µL of each cell culture was lysed in individual wells of a clear, flat-bottomed 96-well plate with 10 µL of PopCulture (Novagen) containing 0.4 mg/mL lysozyme for at least 15 min at room temperature. 20 µL of each cell lysate was transferred to wells containing 100 µL Z-buffer (60 mM Na_2_HPO_4_, 40 mM NaH_2_PO_4_, 10 mM KCl, 1 mM MgSO_4_, 50 mM β-mercaptoethanol at pH 7.0) with 4 mg/mL 2-Nitrophenyl β-D-galactopyranoside (ONPG; Sigma-Aldrich). OD_420_ for each well was read once per min for 1 h at 37°C in a Synergy H1M plate reader (BioTek). β-Galactosidase activity (in arbitrary units [AU]) was calculated as the rate of ONPG conversion (i.e., V_max_, with units of mOD_420_/min) multiplied by a dilution factor of 5 and divided by the OD_600_ of the sample at the time of collection. Previous standard curve analysis determined that 1 AU corresponds to ∼4 Miller units (Camp and Losick 2009).

### Isothermal Titration Calorimetry (ITC)

MdfA, ClpC^N^, or ClpC^N,Q11P^ were purified as described in **Supplemental Materials and Methods** (final buffer conditions: 50 mM Tris pH 8.85, 150 mM NaCl, 50 mM L-Glu, 0.5 mM TCEP). ITC experiments were performed at 25°C using an ITC-200 MicroCal microcalorimeter (GE Healthcare) following a standard procedure (Hands-Taylor et al. 2010). In each titration, 20 injections of MdfA at a concentration of 700 μM were added to ClpC constructs at 70 μM in the reaction cell. Integrated heat data obtained for the titrations, corrected for heats of dilution, were fitted using a nonlinear least-squares minimization algorithm to a theoretical titration curve, using the MicroCal-Origin 7.0 software package.

### Analytical Size Exclusion Chromatography (SEC)

Analytical SEC experiments with MdfA, MecA, and ClpC^DWB^ (purified as described in **Supplemental Materials and Methods**) were performed on a Superose 6 10/300 Increase column (GE Healthcare) at 4°C. Proteins were diluted to 10 µM in column running buffer (50 mM Tris pH 8.0, 300 mM NaCl, 5 mM MgCl_2_, 0.5 mM DTT, 0.5 mM ATP) and incubated 10 min with 5 mM ATP at 37°C as previously described (Kirstein et al. 2006). 1 mL fractions were collected and the proteins were precipitated using trichloroacetic acid (final 10%) and acetone, and analyzed by SDS-PAGE and Coomassie staining. Additional SEC methods are provided in **Supplemental Materials and Methods**.

### ATPase and Protein Degradation Assays

In vitro ATPase assays were performed as previously described (Lanzetta et al. 1979; Turgay et al. 1997). ClpC, ClpC^Q11P^, or ClpC^Q11A^ proteins (1 µM each; purified as described in **Supplemental Materials and Methods**) were mixed with MecA (1 µM) or MdfA (1-15 µM) in ATPase assay buffer (50 mM Tris pH 8.0, 50 mM KCl, 5 mM MgCl_2_). Reactions were started by addition of ATP to a final concentration of 4 mM, at 37°C. Reactions were stopped by adding 10 µL of the reaction mixture to 160 µL staining solution (3.4 mg/L malachite green, 10.5 g/L ammonium molybdate, and 0.1% Triton X-100 in 1 N HCl) in a 96-well plate, followed by the addition of 20 µL 34% (w/v) sodium citrate. Released phosphate was measured by absorbance at 660 nm. ATPase activity was calculated from the linear slope between the 0 min and 5 min time points of the reaction.

### Protein Degradation Assays

Degradation assays were performed in ATPase assay buffer (50 mM Tris pH 8.0, 50 mM KCl, 5 mM MgCl_2_) with an ATP regeneration system (Schlothauer et al. 2003) and all proteins (purified as described in **Supplemental Materials and Methods**) diluted to 1 µM. Reactions were started at 37°C by adding ATP to a final concentration of 4 mM. Samples were taken at indicated time points for analysis by SDS-PAGE followed by Coomassie staining.

### Protein Crystallography

The MdfA/ClpC^N^ complex crystallized at 8.4 mg/mL in 2.0 M (NH_4_)_2_SO_4_, 100 mM NaOAc, pH 4.6. PDB: 5HBN (Trentini et al. 2016) was used as an initial molecular replacement model for ClpC^N^. For further details on screening, data collection, processing and model-building, see **Supplemental Materials and Methods**.

## Author contributions

A.H.C., S.M., R.L.I., N.E., K.T., and I.H. conceived the study. All authors designed and/or performed experiments and analyzed results. A.H.C., S.M., R.L.I., N.E., K.T., and I.H. wrote the original manuscript draft. All authors reviewed and edited the manuscript. A.H.C. and R.L.I. supervised the study. A.H.C., R.L.I., and K.T. acquired funding.

## ACKNOWLEDGEMENTS

This work was funded by grants from the National Institutes of Health (NIH) (DP2 GM105439 to AHC), the Biotechnology and Biological Sciences Research Council (BBSRC) (BB/N006267/1, BB/R006091/1, BB/S006877/1, and BB/X001415/1 to RLI), the Max Planck Society & Deutsche Forschungsgemeinschaft (Tu106/6 and Tu106/8 [SPP1879] to KT), and a Ph.D. fellowship from the Hannover School for Biomolecular Drug Research (HSBDR) (to IH). The authors would like to thank Diamond Light Source for beamtime (proposal mx13597; King’s College London BAG). NMR experiments were performed at the Centre for Biomolecular Spectroscopy, King’s College London, established with a Capital Award from the Wellcome Trust. This work was also supported by the Francis Crick Institute through provision of access to the MRC Biomedical NMR Centre. The Francis Crick Institute received core funding from Cancer Research UK (FC001029), the UK Medical Research Council (FC001029), and the Wellcome Trust (FC001029). The 950 MHz NMR facility at the University of Oxford was funded by the Wellcome Trust Joint Infrastructure Fund and the E. P. Abraham Fund. We also thank R. Wang for constructing pRW3, and R. Stieber, I. Moin, and G. Davis for preliminary data collection. Finally, the authors are deeply grateful for the collegiality of K. Pogliano and members of her research team including J. Lyda, R. Eammon, and J. Lopez-Garrido, as well as for the ongoing and invaluable support of our shared mentor Dr. Richard Losick.

