## Supplemental Materials for "MdfA is a novel ClpC adaptor protein that functions in the developing *Bacillus subtilis* spore"

### SUPPLEMENTAL FIGURES AND LEGENDS

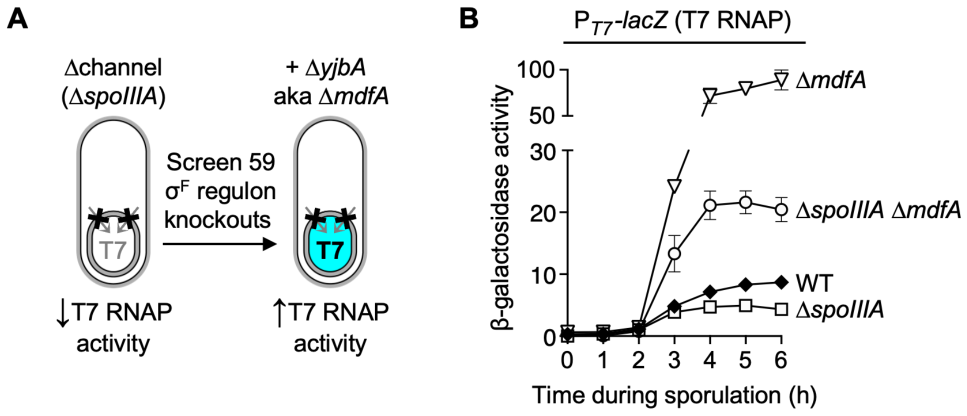


**Figure S1. A genetic screen identifies *yjbA* (renamed here as *mdfA* for *m*etabolic *d*ifferentiation *f*actor *A*) as a candidate gene involved in forespore metabolic shutdown.** **(A)** Graphic depiction of the genetic screen used to identify *mdfA* as a candidate gene involved in forespore metabolic shutdown. This candidate screen was based on two assumptions: first, that such genes would be under the control of $\sigma^F$ and second, that deletion of such genes would cause the forespore to become less dependent upon the SpoIIIAA-AH•SpoIIQ channel for late gene expression. The original screen measured gene expression by a heterologous RNA polymerase (T7 RNA polymerase, from the *E. coli* bacteriophage T7), which has been previously shown to be dependent upon the SpoIIIAA-AH•SpoIIQ channel in the forespore (Camp and Losick 2009) but is unlikely to be the target of any other regulation in *B. subtilis*. Deletions of 59 genes known or predicted to be expressed under the control of $\sigma^F$ (see **Supplemental Table S5**) were transformed individually into a strain expressing T7 RNA polymerase (T7 RNAP) in the forespore and deleted for the channel protein-encoding *spoIIIA* operon (AHB1560). Candidate genes involved in forespore metabolic shutdown were identified as those whose deletions stimulated T7 RNAP activity despite the absence of the SpoIIIAA-AH•SpoIIQ channel.

**(B)** T7 RNAP activity in forespores lacking an active channel is stimulated by Δ*mdfA*, and this stimulation is even more robust in the presence of a functional channel. β-Galactosidase production from the T7 RNAP-dependent $P_{T7}$-*lacZ* reporter was monitored during sporulation of otherwise wild type cells (WT; closed diamonds), cells deleted for *spoIIIA* (Δ*spoIIIA*; open squares), cells deleted for both *spoIIIA* and *mdfA* (Δ*spoIIIA* Δ*mdfA*; open circles), and cells deleted for *mdfA* alone (Δ*mdfA*; open inverted triangles) (strains AHB1449, AHB1560, SMB130, and SMB165, respectively). Error bars indicate ± standard deviations based on two or more independent experiments.

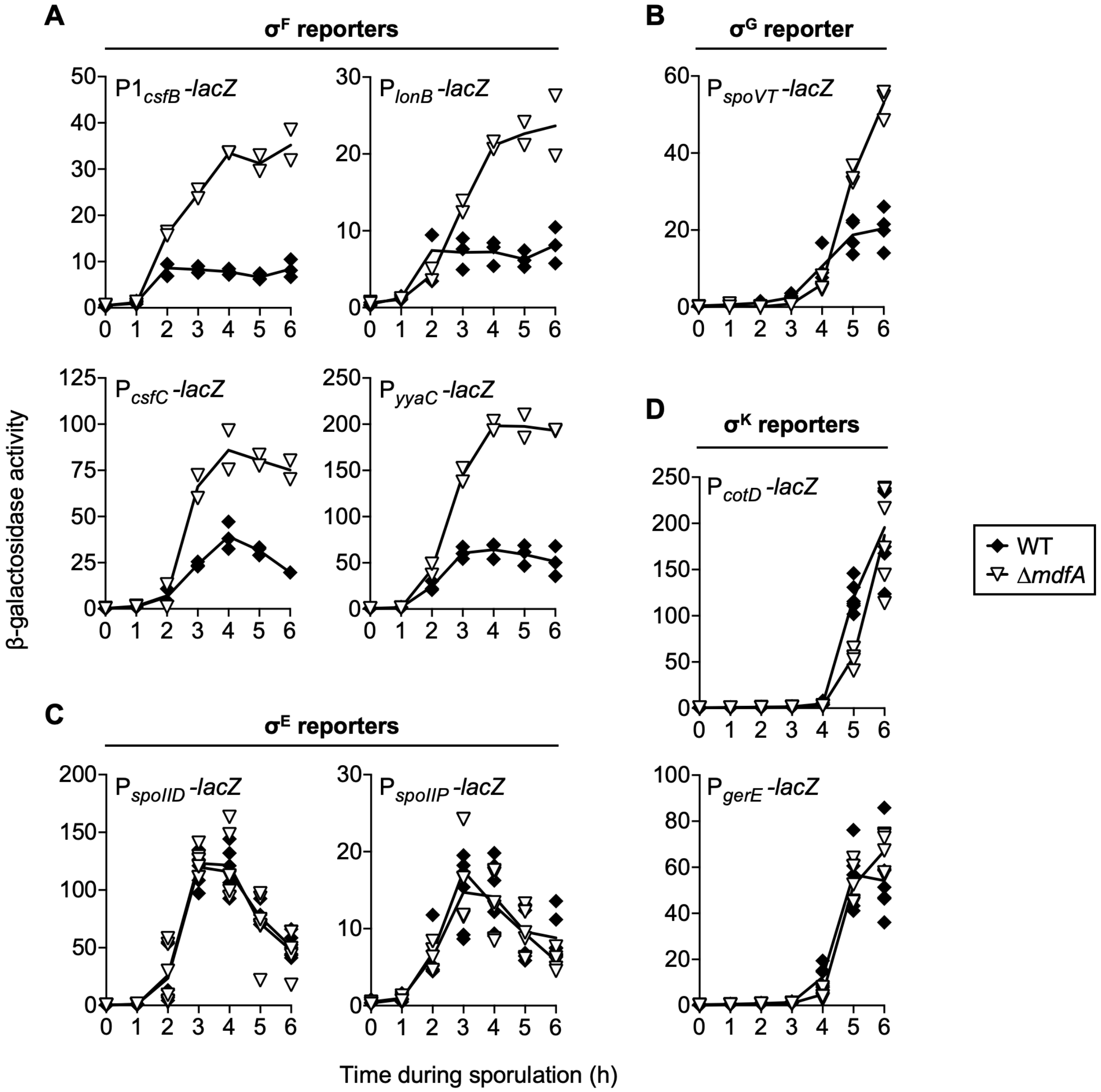


**Figure S2. Δ*mdfA* stimulates forespore ($\sigma^F$ and $\sigma^G$) but not mother cell ($\sigma^E$ and $\sigma^K$) gene expression.**

The expression of **(A)** $\sigma^F$-dependent, **(B)** $\sigma^G$-dependent, **(C)** $\sigma^E$-dependent, and **(D)** $\sigma^E$-dependent *lacZ* reporter genes was monitored during sporulation of otherwise wild type cells (WT; closed diamonds) or cells deleted for the *mdfA* gene (Δ*mdfA*; open inverted triangles). Individual data points are shown (n=2-6) with lines connecting the mean value at each time. Strains (WT, Δ*mdfA*) were as follows: $P1_{csfB}$-*lacZ* (JDC138, SMB302), $P_{lonB}$-*lacZ* (AHB1296, SMB303), $P_{yyaC}$-*lacZ* (AHB1704, SMB304), $P_{csfC}$-*lacZ* (AHB1729, SMB305), $P_{spoVT}$-*lacZ* (BAT87, SMB310), $P_{spoIID}$-*lacZ* (BZ184, CFB153), $P_{spoIIP}$-*lacZ* (PE511, CFB165), $P_{cotD}$-*lacZ* (SMB300, SMB332), $P_{gerE}$-*lacZ* (SMB301, SMB335).

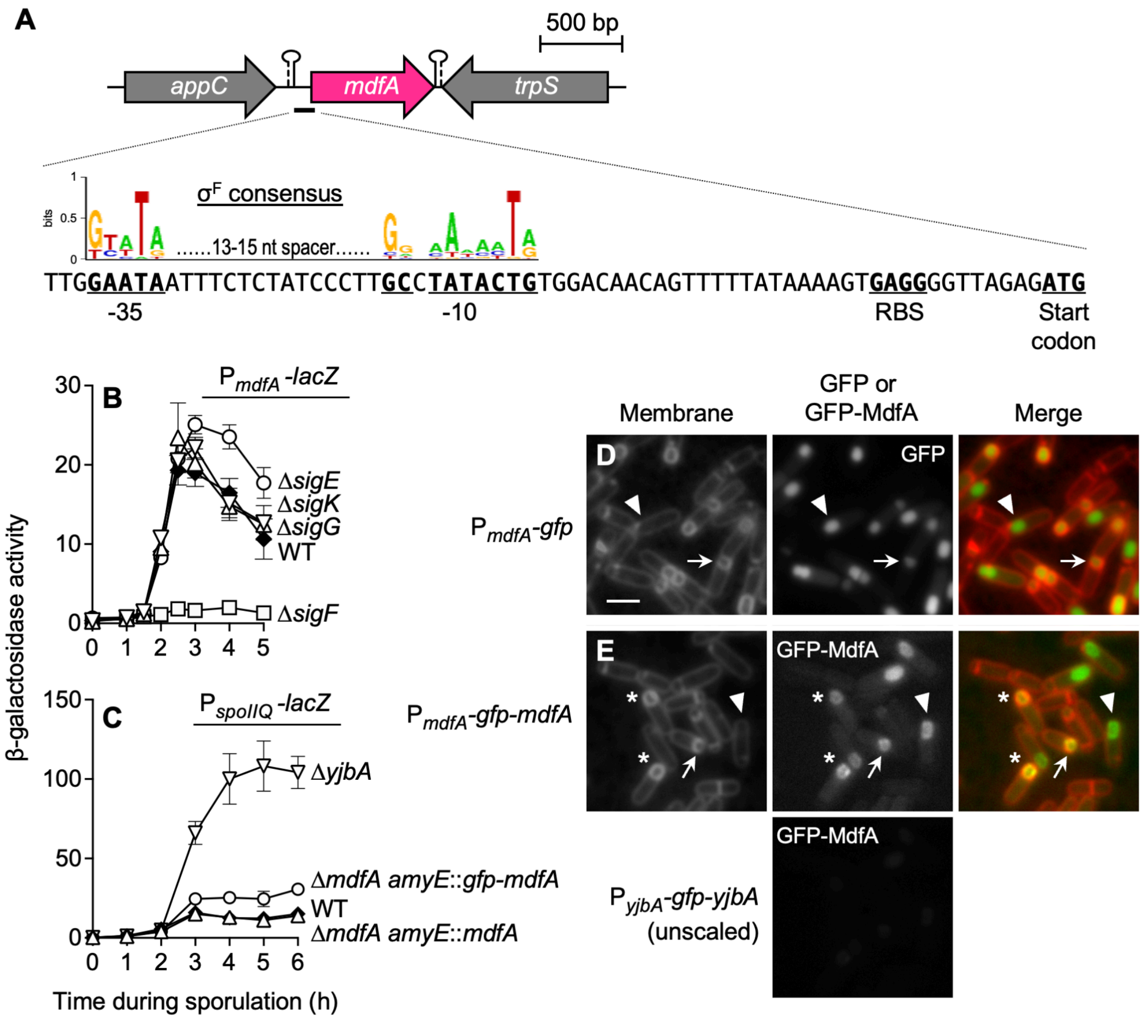


**Figure S3.** ***mdfA* is expressed under σ$^F$ control in the forespore and the MdfA protein localizes to the forespore.**

**(A)** The *mdfA* gene (magenta) depicted within its genomic context. The thick black line immediately upstream of *mdfA* indicates the 122 bp regulatory region, up to and including the *mdfA* start codon, present in the P$_{mdfA}$-*lacZ* and P$_{mdfA}$-*gfp* reporters used in (B), (C), and (D). Zoom-in shows the coding-sequence proximal 71 nt of this region (5' → 3'), which contains putative -10 and -35 promoter elements that are good matches for the σ$^F$ recognition consensus (Sierro et al. 2008). The *mdfA* ATG start codon and putative ribosome-binding site (RBS) are also indicated.

**(B)** *mdfA* is expressed under the control of σ$^F$ during sporulation. Production of β-galactosidase was monitored during sporulation of strains harboring P$_{mdfA}$-*lacZ*, a *lacZ* reporter under the control of *mdfA* transcriptional and translational regulatory sequences shown in (A). Cells were otherwise wild type (WT; closed diamonds) or deleted for the gene encoding σ$^F$ (Δ*sigF*; open squares), σ$^E$ (Δ*sigE*; open circles), σ$^G$ (Δ*sigF*; open triangles), or σ$^K$ (Δ*sigK*; open inverted triangles), (strains SMB266, SMB296, SMB297, SMB299, and SMB298, respectively). Error bars indicate ± standard deviations based on two or more independent experiments.

**(C)** The GFP-MdfA fusion protein used in (E) is almost fully functional. β-Galactosidase production from the $\sigma^F$-dependent $P_{spoIIQ}$-*lacZ* reporter was monitored during sporulation of otherwise wild type cells (WT; closed diamonds), cells deleted for *mdfA* (Δ*mdfA*; open inverted triangles), or cells deleted for *mdfA* and harboring either *mdfA* or *gfp-mdfA* integrated at the *amyE* locus (Δ*mdfA amyE*::*mdfA*; open triangles and Δ*mdfA amyE*::*gfp-mdfA*; open circles, respectively) (strains AHB1841, CFB530, CFB541, and CFB545, respectively). Error bars indicate ± standard deviations based on two independent experiments.

**(D,E)** *mdfA* expression is confined to the forespore and GFP-MdfA localizes to the forespore during sporulation. Cells expressing **(D)** GFP alone or **(E)** a functional GFP-MdfA fusion protein under the control of *mdfA* transcriptional and translational regulatory sequences ($P_{mdfA}$-*gfp* or $P_{mdfA}$-*gfp-mdfA*, respectively) were observed by fluorescence microscopy at sporulation hour 3 (strains SMB409 and SMB434, respectively). GFP fluorescence is shown in grayscale (GFP or GFP-MdfA) or false-colored green (Merge). The panel labeled "$P_{mdfA}$-*gfp-mdfA* (unscaled)" shows the fluorescence intensity of GFP-MdfA when settings were identical to those of GFP alone. Membrane fluorescence from the dye FM 4-64 is shown in grayscale (Membrane) or false-colored red (Merge). White arrows indicate forespores at intermediate stages of engulfment, while the white arrowheads indicate forespores that have been fully engulfed. Note that membranes surrounding engulfed forespores are not stained due to membrane impermeability of the FM 4-64 dye. Asterisks indicate two cells (among others) that display peripheral membrane localization. Scale bar = 2 μm.

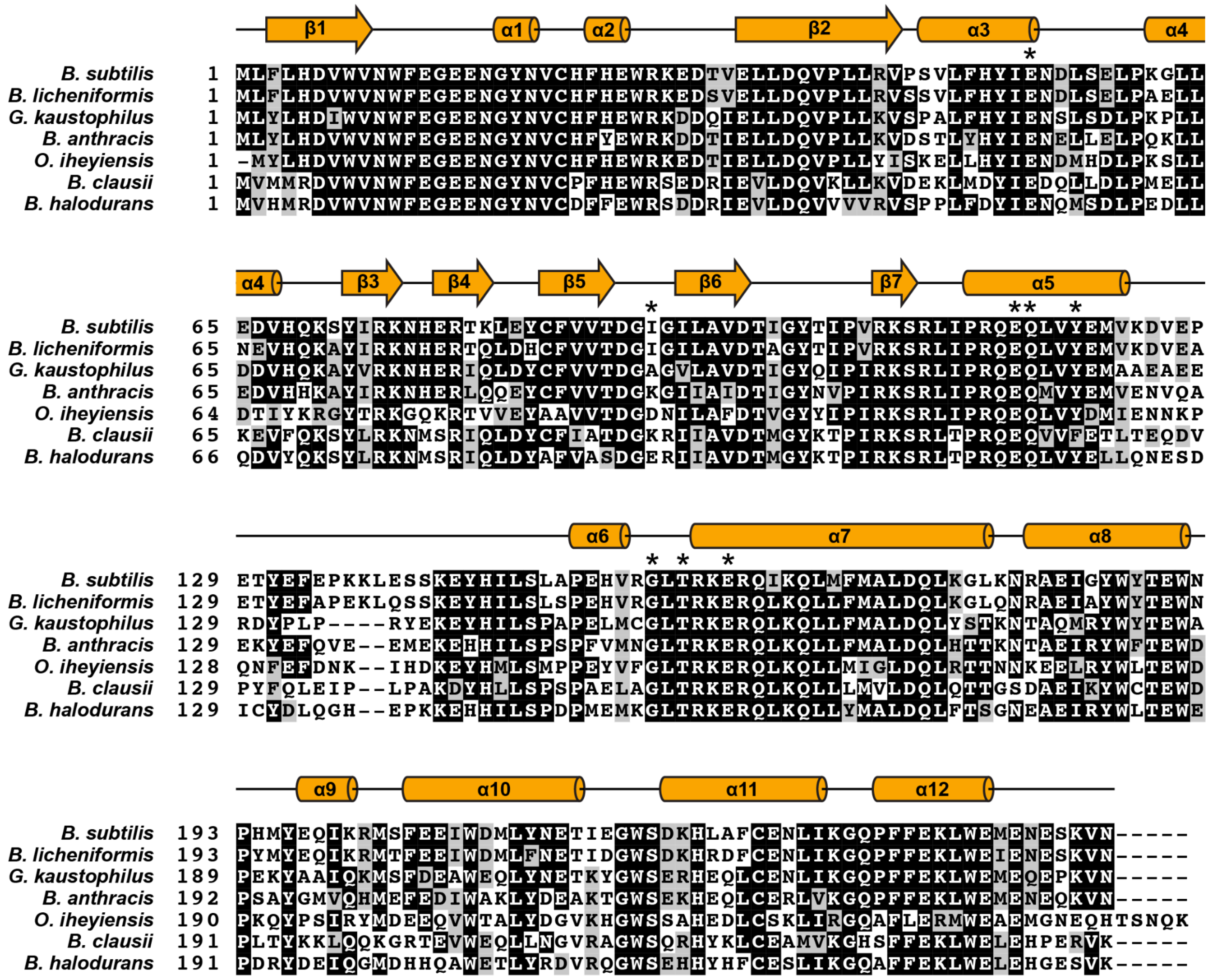


**Figure S4. The MdfA protein is conserved among the *Bacillaceae*.**
A multiple sequence alignment of the MdfA proteins from representative members of the *Bacillaceae* family was generated with Clustal Omega (Sievers et al. 2011) and shaded with pyBoxshade. Amino acid residues displaying >60% identity or similarity are shaded black or gray, respectively. Asterisks (*) indicate residues making polar contacts with the ClpC N-domain in the MdfA-ClpC$^{N}$ co-crystal structure reported here. Secondary structures (β sheets and α helices) from the MdfA structure reported here are shown in cartoon form above the alignment. The following MdfA reference sequences were selected as representative orthologs (GenBank accession numbers are given in parentheses): *Bacillus subtilis* (NP_391536.1), *Bacillus licheniformis* (WP_003185941.1), *Geobacter kaustophilus* (WP_011232806.1), *Bacillus anthracis* (NP_847683.1), *Oceanobacillus iheyensis* (WP_011067358.1), *Bacillus clausii* (WP_011248679.1), and *Bacillus halodurans* (WP_010899871.1).

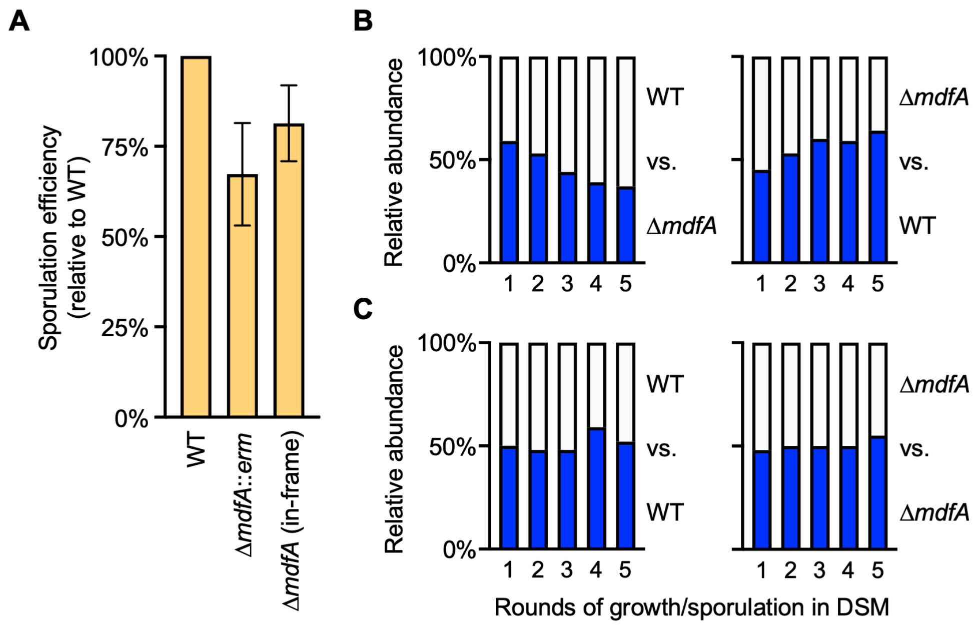


**Figure S5. *mdfA* is required for maximal sporulation efficiency**

**(A)** Cells lacking *mdfA* sporulate with a lower efficiency than wild type cells in single-round sporulation assays. Wild type cells (WT) or cells harboring a Δ*mdfA*::*erm* or in-frame Δ*mdfA* deletion (strains PY79, SMB251, and SMB431, respectively) were induced to sporulate by nutrient exhaustion in DSM for 24 h. Sporulation efficiency was calculated as the number of heat-resistant spores produced by each mutant normalized to those produced by wild-type, which was set to 100%. Data are presented as the mean ± standard deviation (n = 3)

**(B)** Cells harboring an in-frame Δ*mdfA* deletion are at a competitive disadvantage during successive rounds of growth and sporulation in co-culture with wild type (WT) cells. WT and Δ*mdfA* cells were inoculated into DSM at an ~1:1 ratio. After 24 h of growth and sporulation, the ratio of heat-resistant spores of each genotype was determined by the presence (blue) or absence (white) of an IPTG-inducible *lacZ* reporter gene. Surviving spores were then back-diluted into fresh medium for another round of growth and sporulation. The relative abundance of each cell type is graphed as the percentage of the total colony forming units (e.g. surviving spores) after each 24 h round of co-culture and heat treatment. The competitive disadvantage of Δ*mdfA* cells was apparent regardless of which genotype harbored the identifying *lacZ* reporter (left vs. right).

**(C)** Control competitions between strains with identical genotypes (WT vs. WT [left] and Δ*mdfA* vs. Δ*mdfA* [right]) further verified that the identifying IPTG-inducible *lacZ* reporter did not confer a (dis)advantage to either strain. The data presented in each of the four graphs for (B) and (C) are averages of triplicate experiments. Strains and relevant genotypes are as follows: CFB123 (WT *lacZ*$^{+}$), CFB125 (WT [no *lacZ*]), CFB602 (Δ*mdfA lacZ*$^{+}$), and CFB604 (Δ*mdfA* [no *lacZ*]).

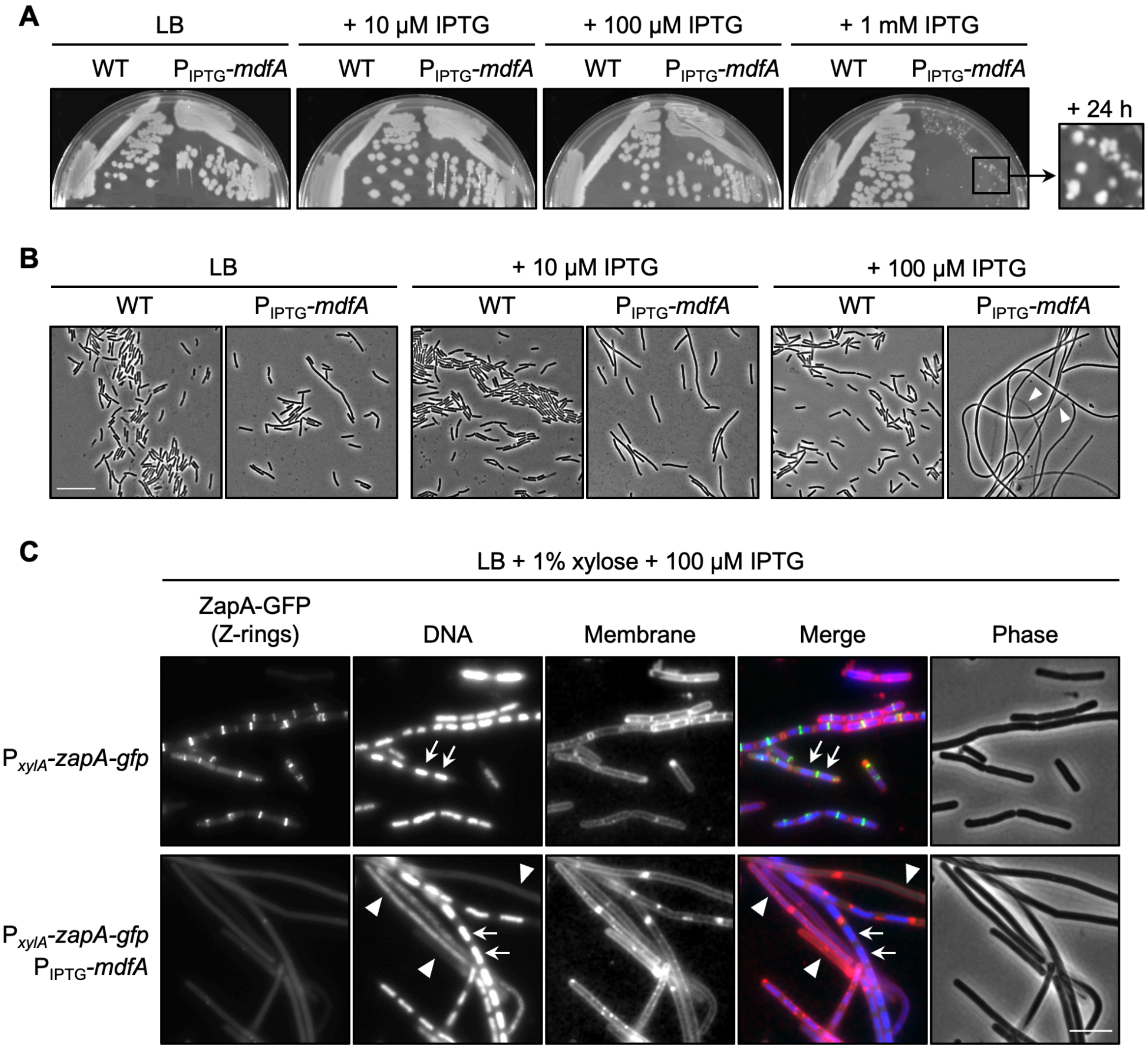


**Figure S6.** ***mdfA*** **expression during vegetative growth causes cell filamentation and lysis.**
**(A)** Expression of *mdfA* during vegetative growth on solid growth medium is toxic at high inducer concentrations. Wild type cells (WT; strain PY79) and cells harboring an engineered IPTG-inducible *mdfA* gene construct ($P_{IPTG}$-*mdfA*; strain CFB189) were grown on LB agar plates without or with 10 µM, 100 µM, or 1 mM IPTG. Plates were imaged after 24 h of growth at 37˚C. Inset shows the spontaneous appearance and robust growth of $P_{IPTG}$-*mdfA* suppressor mutant colonies on LB plates with 1 mM IPTG after an additional 24 h of growth at 37˚C.
**(B)** Expression of *mdfA* during vegetative growth causes cell filamentation even at low inducer concentrations, in a dose-dependent manner. WT and $P_{IPTG}$-*mdfA* cells (strains PY79 and CFB189, respectively) were grown for ~24 h at 37˚C on LB agar plates without or with 10 µM or 100 µM IPTG. Colonies were picked, diluted, and visualized by phase microscopy. Arrowheads indicate cell filaments that have undergone lysis. Scale bar, 10 µm.

**(C)** FtsZ rings fail to form in vegetative cells expressing *mdfA*, although chromosomes are often properly partitioned. WT and $P_{IPTG}$-*mdfA* cells with a $P_{xyl}$*-GFP-zapA* reporter (strains CFB525 and CFB549, respectively) were grown for ~24 h at 37˚C on LB agar plates with 100 µM IPTG and 1% xylose (to induce expression of GFP-ZapA). Colonies were picked, diluted, stained with the membrane dye FM 4-64 and DNA stain DAPI, and visualized by phase and fluorescence microscopy. GFP fluorescence, DAPI, and FM 4-64 staining are shown in grayscale (ZapA-GFP, DNA, and Membrane, respectively) or false-colored (Merge; green, blue, and red, respectively). Scale bar, 10 µm.

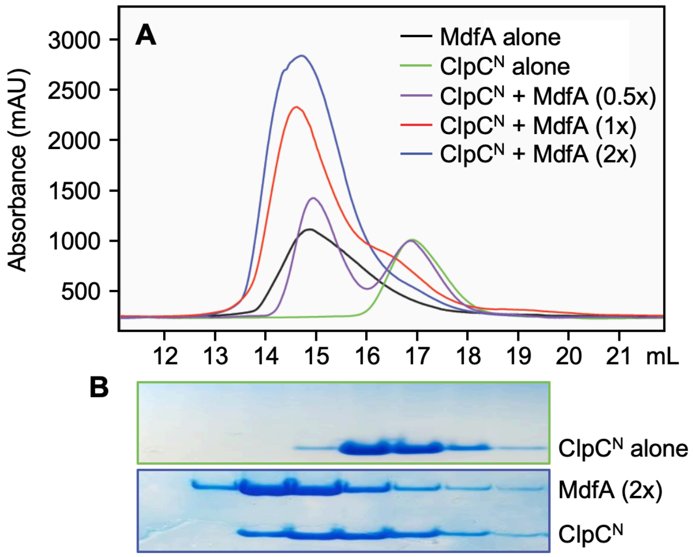


**Figure S7. MdfA forms a complex with ClpC$^{N}$ as detected by size exclusion chromatography.**

MdfA and ClpC$^{N}$ alone (at 266 µM each) or in various molar ratios (as indicated, with ClpC held constant at 266 µM) were separated by size exclusion chromatography and eluted protein was detected **(A)** by absorbance at 230 nm (to account for the widely differing extinction coefficients of the two proteins) or **(B)** by SDS-PAGE.

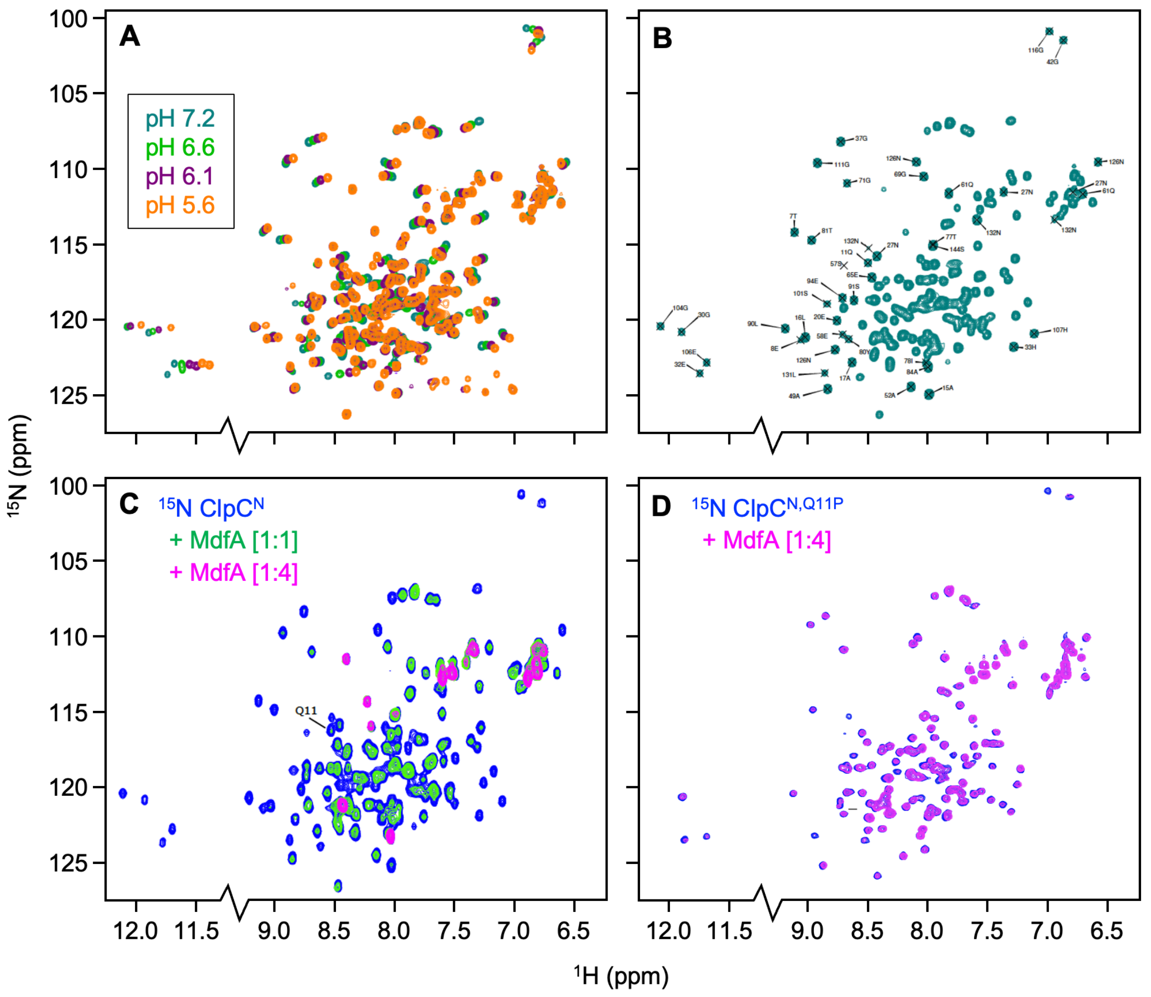


**Figure S8. MdfA forms a complex with ClpC$^N$ in a Q11-dependent manner, as detected by NMR titration experiments.**

**(A,B)** ClpC$^N$ NMR peak assignments as determined by pH titration. To compare our $^{15}$N-$^1$H HSQC spectra of free ClpC$^N$ obtained at pH 7.2 (teal) with a previously published assignment obtained at pH 5.5 (Biological Magnetic Resonance Data Bank [BMRB Entry: 15383]) (Kojetin et al. 2007, 2009), we performed a pH titration (pH 6.6 [green], pH 6.1 [purple], and pH 5.6 [orange]) to follow individual peak positions, shown in **(A)**. The resulting peak assignments are shown in **(B)**.

**(C,D)** NMR titration experiments indicate that MdfA and $^{15}$N-labeled ClpC$^N$ but not ClpC$^{N,Q11P}$ (ClpC$^N$ harboring the Q11P substitution) interact *in vitro*. $^{15}$N-$^1$H HSQC spectra are shown for **(C)** ClpC$^N$ or **(D)** ClpC$^{N,Q11P}$ either alone (blue) or in the presence of MdfA at a 1:1 molar ratio (green) or 1:4 molar ratio (magenta). The position of residue Q11 in the ClpC$^N$ HSQC spectra is labeled (Q11).

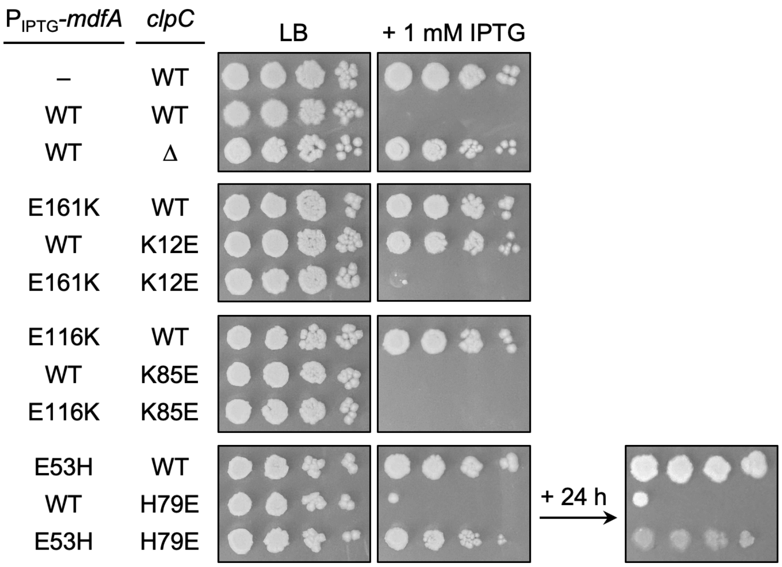


**Figure S9.** Mutational analysis of the MdfA-E161/ClpC-K12, MdfA-E116/ClpC-K85, and MdfA-E53/ClpC-H79 electrostatic interactions in MdfA toxicity during vegetative growth. Cells harboring an engineered IPTG-inducible *mdfA* gene construct ($P_{IPTG}$-*mdfA*) with either wild type (WT) or mutant *mdfA* alleles (E161K, E116K, or E53H), and harboring either wild type (WT), deletion (Δ), or other mutant alleles (K12E, K85E, or H79E) of *clpC* at its native locus were grown for 6 hours in LB media at 37°C, subjected to 10-fold serial dilutions ($10^{-2}$-$10^{-5}$, left to right), and spotted using a sterile 48 pin custom-made replicator onto LB plates with or without 1 mM IPTG. A wild type strain without $P_{IPTG}$-*mdfA* (–) was also included as a control. Note that the native *mdfA* gene was left unaltered in these strains. Plates were grown at 25°C and were imaged after 48 h and, in the case of the bottom far right panel, 72 h. Note that the $P_{IPTG}$-$mdfA^{E53H}$ $clpC^{H79E}$ strain shows a small colony phenotype in the presence of inducer and evidence of cell lysis after another 24 h of growth. Strains were as follows (listed as in the figure from top to bottom): PY79, CFB189, CFB241, CSB8, RFB21, AHB6213, XWB12, XWB50, XWB58, SFB8, SOB18, AHB6215.

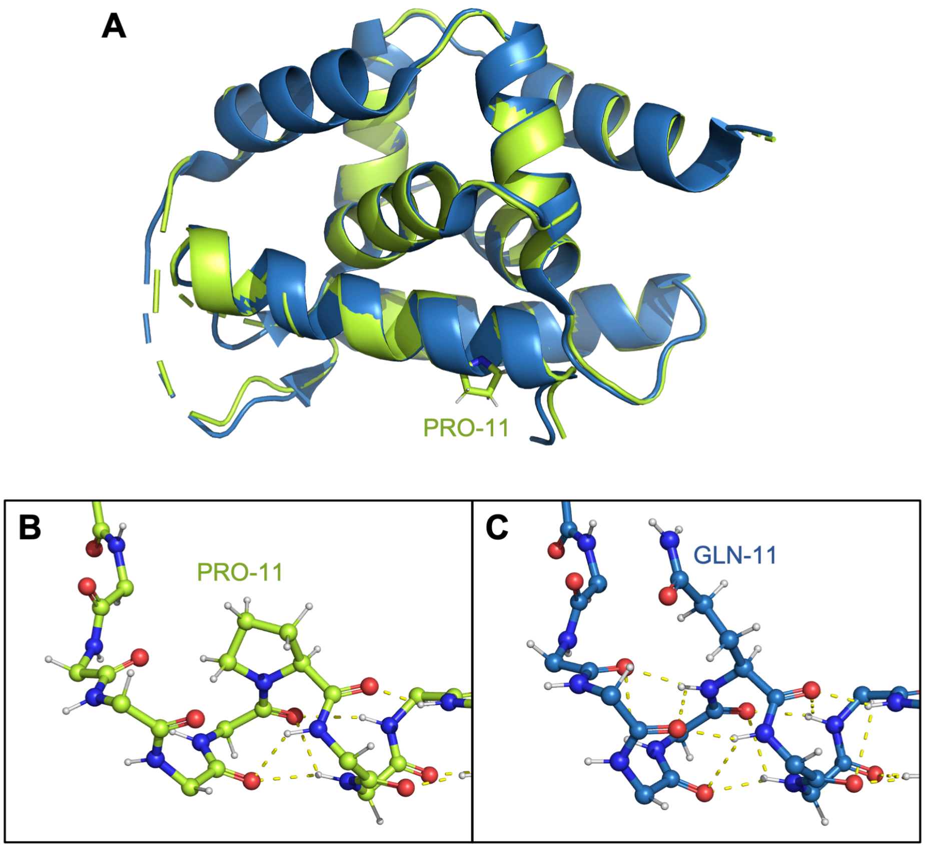


**Figure S10. X-ray crystal structure of ClpC$^{N,Q11P}$**

**(A)** Overlay of the ClpC$^{1\text{-}150,Q11P}$ structure (lime green) with wild type ClpC$^{N}$ (blue; PDB: 2Y1Q)**.** **(B)** Hydrogen bonding network in helix 1 of ClpC$^{N,Q11P}$ structure. **(C)** Equivalent hydrogen bonding network around Q11 in wild-type ClpC$^{N}$.

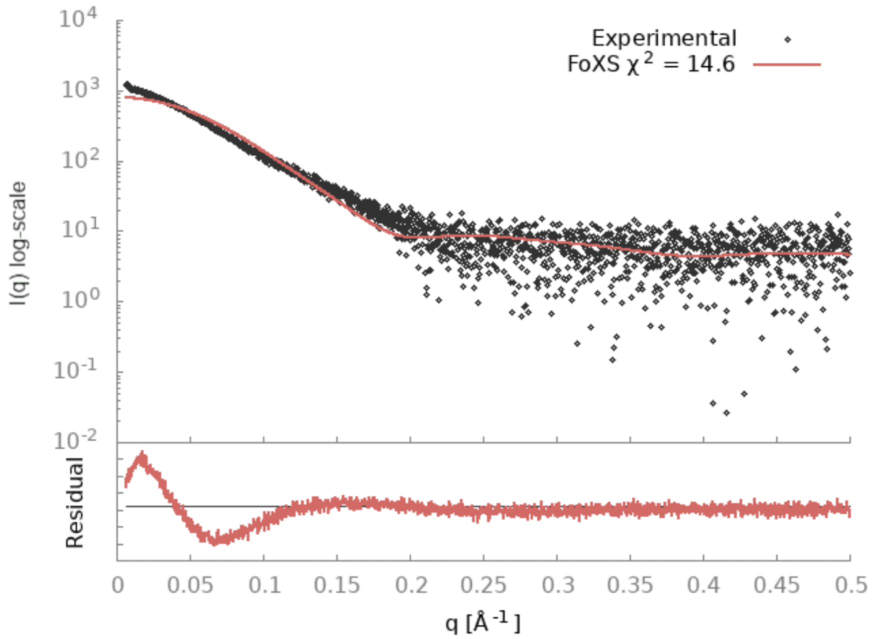


**Figure S11. SAXS data for isolated MdfA indicate greater flexibility than MdfA displays in crystal complex with ClpC$^{N}$**

The FoXs data server (Schneidman-Duhovny et al. 2013, 2016) was used to attempt to fit the MdfA monomer unit (excised from the crystal structure) to the scattering data. The result shown is visually not a good fit to the data, a fact confirmed by the high chi-squared measure. In this fit large errors are present in the low q range of the data corresponding to the large scale geometry of the protein. The data have a significantly linear downward slope up to q=0.2 indicative of an elongated structure in contrast to the model PDB curve which has a decreasing slope more indicative of globular shapes. These data indicate a strong likelihood that the structure adopts a more elongated shape alone in solution than it does in the complex crystal structure.

### SUPPLEMENTAL MATERIALS AND METHODS

#### *B. subtilis* strain construction

All *B. subtilis* strains used in this study were isogenic with the laboratory strain PY79 (Youngman et al. 1984). Strains were typically propagated in lysogeny broth (LB), in liquid culture or on solid plates with 1.6% agar. When appropriate, antibiotics were included as follows, unless otherwise indicated: chloramphenicol (5 µg/mL), erythromycin plus lincomycin (1 µg/mL and 25 µg/mL, respectively), spectinomycin (100 µg/mL), and kanamycin (5 µg/mL). Antibiotic resistance genes are referred to as follows: *cat* (chloramphenicol), *erm* (erythromycin plus lincomycin), *spc* (spectinomycin), and *kan* (kanamycin). Derivatives of PY79 were generated by transformation with chromosomal DNA from *B. subtilis*, plasmid DNA, or DNA fragments generated by PCR and/or isothermal assembly. Competent *B. subtilis* cells were prepared as previously described (Wilson and Bott 1968). Chromosomal integration into the *amyE* locus was confirmed by loss of α-amylase activity on LB agar plates with 1% starch. Insertion into the *"*alternative *amyE"* sites at *ywrK* and *ylnF* were as previously described (Camp and Losick 2008, 2009). The full genotypes of *B. subtilis* strains used in this study are given in **Supplemental Table S1**. Plasmids used to construct strains are provided in **Supplemental Table S2**. Details of plasmid design and construction can be found in the "Plasmid construction" section, below.

**Deletion mutants.** The Δ*sigF*::*erm* and Δ*sigE*::*erm* deletions were from strains RL1275 and RL1061 (Eichenberger et al. 2001), respectively. The Δ*sigG*::*kan* and Δ*spoIIIAA-AH*::*erm*::*phleo* deletions were from strains AHB98 (Camp and Losick 2008) and AHB1225 (Camp and Losick 2009), respectively. The Δ*spoIVCB*::*erm* deletion, which removes part of the composite gene for *sigK*, was from laboratory stock strain AHB51, built by transforming Δ*spoIVCB*::*erm* from MO1027 (gift of P. Stragier) into PY79. All other deletion mutants were from the *B. subtilis* "BKE" Knockout Collection (Koo et al. 2017), distributed by the *Bacillus* Genetic Stock Center (BGSC). The BKE knockouts used in the metabolic differentiation factor screen are listed in **Supplemental Table S5**, including Δ*mdfA(yjbA)*::*erm* (BKE11410). Other BKE knockouts used in this study were Δ*clpC*::*erm* (BKE00860), Δ*clpP*::*erm* (BKE34540), and Δ*yacL*::*erm* (BKE00890). Given that the BKE knockouts were constructed in the auxotrophic *B. subtilis* strain 168, these gene deletions were moved into PY79 or PY79-derived strains prior to analysis. The Δ*mdfA*::*erm*::*spc* deletion was constructed by transforming SMB251 (Δ*mdfA*::*erm* from BKE11410 in PY79), with plasmid pEr::Sp (Steinmetz and Richter 1994), yielding the erythromycin/lincomycin-sensitive, spectinomycin-resistant strain CFB313. The in-frame Δ*mdfA* knockout (SMB432) was constructed by removing the erythromycin resistance cassette from SMB251 using the temperature sensitive, Cre recombinase expressing plasmid pDR244, as previously described (Koo et al. 2017).

**Unmarked *clpC* and *mdfA* mutants.** The $clpC^{Q11P}$ mutant originally isolated as a suppressor of *mdfA* toxicity during vegetative growth (see "Isolation and identification of *mdfA* extragenic suppressor mutants" section, below) was, in some cases, moved into new strains through linkage to an adjacent antibiotic resistance gene (*erm*), inserted in the non-essential *yacL* gene. Transformants were screened by PCR-amplification and sequencing of their *clpC* loci to identify those that had acquired the *Q11P* mutation. The Δ*yacL*::*erm* deletion alone was confirmed to have no effect in all relevant assays. Otherwise, unmarked *clpC* mutants (*clpC*$^{Q11P}$, *clpC*$^{Q11A}$, *clpC*$^{K12E}$, *clpC*$^{H79E}$*, and clpC*$^{K85E}$) were constructed from scratch with the temperature-sensitive plasmids pSF15 (*clpC*$^{Q11P}$ in pMiniMAD2), pSF19 (*clpC*$^{Q11A}$ in pMiniMAD2), pRF6 (*clpC*$^{K12E}$ in pMiniMAD2), pSO3 (*clpC*$^{H79E}$ in pMiniMAD2), and pXW22 (*clpC*$^{K85E}$ in pMiniMAD2) respectively, using a loop-in/loop-out method (Patrick and Kearns 2008). Single-crossover integration of the plasmids at the *clpC* locus was selected at the restrictive temperature (37°C) in the presence of antibiotic (erythromycin plus lincomycin). Cells were then grown at the permissive temperature (25°C) in the absence of antibiotic to evict the integrated plasmid. Individual erythromycin/lincomycin sensitive colonies were screened by PCR-amplification and sequencing of their *clpC* loci to identify those that had acquired the desired mutations.

Mutations in the native *mdfA* gene (*mdfA*$^{E53H}$, *mdfA*$^{E116K}$, and *mdfA*$^{E161K}$) were constructed using CRISPR-Cas9 gene editing as described by (Sachla et al. 2021). Plasmids pAV3 (*mdfA*$^{E53H}$ repair template in pAJS23), pRW3 (*mdfA*$^{E116K}$ repair template in pAJS23), and pSY3 (*mdfA*$^{E161K}$ repair template in pAJS23), were transformed into strain SMB189 (Δ*mdfA*::*erm amyE*::P$_{spoIIQ}$-*lacZ cat*) with selection at 25°C (permissive for plasmid replication) on LB agar plates supplemented with 15 µg/mL kanamycin (to select for the plasmid) and 0.2% mannose (to induce expression of Cas9). Cells were then grown at the non-permissive temperature (45°C) in the absence of antibiotic to evict the plasmid. Individual colonies that lost erythromycin/lincomycin resistance (indicating successful gene editing) and kanamycin resistance (indicating plasmid loss) were screened by PCR-amplification and sequencing of their *clpC* loci to confirm acquisition of the desired mutations.

**T7 RNA polymerase (T7 RNAP) expression construct.** The P$_{spoIIQ}$-*T7RNAP* expression construct, integrated at the *ylnF* alternative *amyE* locus has been previously described (Camp and Losick 2009).

***mdfA* complementation and overexpression constructs.** The *P*$_{mdfA}$-*mdfA* complementation constructs, integrated either at *amyE* or the *ywrK* alternative *amyE* locus, were built for this study (see plasmid pSM1). The IPTG-inducible *P*$_{hyperspank}$-*mdfA* overexpression constructs, integrated at either *amyE* or *thrC*, were built for this study (see plasmids pSM26 and pSM30, respectively).

***lacZ* reporter constructs.** The *lacZ* reporter constructs used in this study were integrated at the *amyE* locus or one of the alternative *amyE* loci at *ywrK* or *ylnF*. The *P*$_{spoIIQ}$-*lacZ*, *P*$_{sspB}$-*lacZ*, and *P*$_{T7}$-*lacZ* reporters were previously described (Camp and Losick 2009), as were P$_{yyaC}$-*lacZ*, P1$_{csfB}$-*lacZ*, and P$_{lonB}$-*lacZ* (Flanagan et al. 2016). The P$_{spoVT}$-*lacZ* reporter fusion was from strain BAT87 (gift of B. Traag). The P$_{csfC}$-*lacZ* reporter fusion was obtained from laboratory stock strain AHB1729, which was built by moving the P$_{csfC}$-*lacZ* reporter from strain RL1310 (gift of R. Losick) into PY79. We note here that the $\sigma^F$-dependent *csfC* promoter, which is not annotated in most genome databases, is located in the intergenic region between *murAA* and *spoIID* and is oriented such that it is predicted to transcribe the non-template strand of *murAA*; it is not thought to encode a protein (Decatur and Losick 1996). The P$_{spoIID}$-*lacZ* reporter was obtained from strain BZ184 (gift of R. Losick), which was built by moving P$_{spoIID}$-*lacZ* from MO367 (Stragier et al. 1988) into PY79. The P$_{spoIIP}$-*lacZ* reporter was obtained from strain PE511 (Eichenberger et al. 2004; gift of R. Losick), which was built by moving P$_{spoIID}$-*lacZ* from

MO1533 (Frandsen and Stragier 1995) into PY79. The remaining reporters (*$P_{cotD}$-lacZ*, *$P_{gerE}$-lacZ*, and *$P_{mdfA}$-lacZ*) were built for this study (see plasmids pAH412, pAH413, and pSM12, respectively). The IPTG-inducible $P_{hyperspank}$-*lacZ* reporter (used to distinguish strains during competition assays) was from RL2508 (gift of R. Losick). The control "empty" $P_{hyperspank}$-[no *lacZ*] construct was from pDR111 (amyE::$P_{hyperspank}$- *lacI spc*; gift of D. Rudner).

***gfp* reporter/fusion constructs.** The *gfp* reporter/fusion constructs used in this study were integrated at the *amyE* locus. The *$P_{mdfA}$-gfp* reporter was built for this study (see plasmid pSM59). The construct expressing *gfp-mdfA* under the native *mdfA* promoter ($P_{mdfA}$-*gfp-mdfA*) was also built for this study (see plasmid pSM60). The xylose-inducible *gfp-zapA* construct was obtained from plasmid pFG28, which was previously described (Gueiros-Filho and Losick 2002).

**Plasmid construction**

Plasmids used in this study are listed in **Supplemental Table S2**. The sequences of primers used in plasmid construction are given in **Supplemental Table S3**. Chromosomal DNA from PY79 served as a template for PCR, unless otherwise noted. Information on synthetic gene fragments (gBlocks, IDT Inc.) used in plasmid construction is given in **Supplemental Table S4**. Plasmids were constructed by isothermal assembly using the Gibson Assembly Master Mix (New England Biolabs) or NEBuilder HiFi DNA Assembly Master Mix (New England Biolabs) and were propagated in the *E. coli* strain NEB 5-alpha (New England Biolabs), unless otherwise noted. All plasmids were verified by sequencing. Details of plasmid construction are provided below.

**pAH412** (*amyE*::$P_{cotD}$-*lacZ cat, amp*) and **pAH413** (amyE::$P_{gerE}$-*lacZ cat, amp*) were designed to integrate a *lacZ* reporter gene fused to the promoter region of *cotD* ($P_{cotD}$) or *gerE* ($P_{gerE}$), respectively, at the *B. subtilis amyE* locus. The *cotD* or *gerE* upstream regulatory sequences, including the promoter, 5' leader sequence, RBS, and start codon, were PCR-amplified with primer pairs AH234/AH383 (for $P_{cotD}$) or AH62/AH384 (for $P_{gerE}$), digested with EcoRI/HindIII, and ligated into EcoRI/HindIII/CIP-treated pAH124 (*amyE*::*lacZ cat*, *amp*) (Camp and Losick 2009).

**pSF15** (*$clpC^{Q11P}$* in pMiniMAD2), **pSF19** (*$clpC^{Q11A}$* in pMiniMAD2), **pRF6** (*$clpC^{K12E}$* in pMiniMAD2), **pSO3** (*$clpC^{H79E}$* in pMiniMAD2), and **pXW22** (*$clpC^{K85E}$* in pMiniMAD2) were designed to insert the *Q11P*, *Q11A*, *K12E*, *H79E*, and *K85E* mutations, respectively, at the native *clpC* locus using a loop-in/loop-out strategy. First, a ~2.2 kb wild type DNA fragment centered on the 5' end of the *clpC* gene encoding the N-domain was PCR-amplified with primer pair oSF5/oSF6 and assembled into EcoRI/BamHI-digested pMiniMAD2 (*ori[ts] erm amp*) (Patrick and Kearns 2008), yielding **pSF14** (*clpC* in pMiniMAD2). Next, pSF14 was subject to site-directed mutagenesis (Q5 Site-Directed Mutagenesis Kit, New England Biolabs) using primer pairs oSF7/oSF9 (*Q11P*), oSF8/oSF9 (*Q11A*), oCS8/oCS9 (*K12E*), oSF11/oSF12 (*H79E*), or oXW6/oXW7 (*K85E*). Plasmids were transformed into and harvested from the *$recA^+$* *E. coli* strain NEB Turbo (New England Biolabs) to produce plasmid concatamers for strain construction.

**pAV3** (*$mdfA^{E53H}$* repair template in pAJS23), **pRW3** (*$mdfA^{E116K}$* repair template in pAJS23), and **pSY3** (*$mdfA^{E161K}$* repair template in pAJS23) were designed to insert the *E53H*, *E116K*, and *E161K* mutations, respectively, at the native *mdfA* locus using CRISPR-Cas9 gene editing as described by (Sachla et al. 2021). First, a ~2.9 kb wild type *mdfA* repair template, harboring *mdfA* coding and flanking sequences, was inserted between the SfiI sites of pAJS23 (*ori[ts] cas9 erm-gRNA kan*) (Sachla et al. 2021) using FastCloning (Li et al. 2011). Both pAJS23 and the *mdfA* repair template were amplified by PCR (using primer pairs oSY1/oSY2 and oSY3/oSY4, respectively), mixed together, digested with DpnI to remove pAJS23 PCR template plasmid, and transformed into the *E. coli* strain NEB 10-beta (New England Biolabs) for in vivo assembly, yielding **pSY1** (*mdfA* repair template in pAJS23). Next, pSY1 was subject to site-directed mutagenesis using the method described by Liu and Naismith (Liu and Naismith 2008) with primer pairs oAA2/oAA3 (*E53H*), oRW4/oRW5 (*E116K*), or oSY8/oSY9 (*E161K*). PCR products were DpnI digested to remove pSY1 PCR template plasmid and transformed into the *E. coli* strain NEB 10-beta (New England Biolabs) for in vivo assembly.

**pSM1** (*amyE*::$P_{mdfA}$-*mdfA kan*, *amp*) was designed to integrate *mdfA* under the control of its own promoter ($P_{mdfA}$) at the *B. subtilis amyE* locus. To construct this plasmid, $P_{mdfA}$-*mdfA* was PCR-amplified with primer pair pr*yjbA*FC/pr*yjbA*RC and assembled into EcoRI/BamHI-digested pER82 (*amyE*::*kan*, *amp*) (Driks et al. 1994).

**pSM12** (*amyE*::$P_{mdfA}$-*lacZ cat, amp*) was designed to integrate a *lacZ* reporter gene fused to the promoter region of *mdfA* ($P_{mdfA}$) at the *B. subtilis amyE* locus. To construct this plasmid, the *mdfA* upstream regulatory sequences, including the putative promoter, 5' leader sequence, RBS, and start codon, were PCR-amplified with primer pair pr*yjbAA*PF/pr*yjbA*PR and assembled into EcoRI/HindIII-digested pAH124 (*amyE*::*lacZ cat*, *amp*) (Camp and Losick 2009).

**pSM26** (*amyE*::$P_{hyperspank}$-*mdfA lacI spc*, *amp*), **pCS2** (*amyE*::$P_{hyperspank}$-*mdfA*$^{E161K}$ *lacI spc*, *amp*), **pSF2** (*amyE*::$P_{hyperspank}$-*mdfA*$^{E53H}$ *lacI spc*, *amp*), and **pXW7** (*amyE*::$P_{hyperspank}$-*mdfA*$^{E116K}$ *lacI spc*, *amp*) were designed to integrate an IPTG-inducible *mdfA* construct (wild type, E161K, E53H, or E116K allele, respectively) at the *B. subtilis amyE* locus. To construct pSM26, the *mdfA* coding sequence was PCR-amplified using primer pair prSM27/prSM28 by a strategy that incorporated an optimized RBS at an ideal spacing from the *mdfA* start codon. This fragment was assembled into SalI/SphI-digested pDR111 (*amyE*::$P_{hyperspank}$- *lacI spc*, *amp*) (gift of D. Rudner). To construct pCS2, pSF2, and pXW7, pSM26 was subject to site directed mutagenesis (Q5 Site-Directed Mutagenesis Kit, New England Biolabs) using primer pairs oCS3/oCS4 (*E161K*), oSF3/oSF4 (*E53H*), and oAK3/oAK4 (*E116K*), respectively.

**pSM30** (*thrC*::$P_{hyperspank}$-*mdfA lacI erm*, *amp*) was designed to integrate an IPTG-inducible *mdfA* construct at the *B. subtilis thrC* locus. To construct this plasmid, the BamHI/EcoRI fragment harboring $P_{hyperspank}$-*mdfA* and *lacI* from pSM26 was subcloned by traditional ligation into BamHI/EcoRI-digested pDG1664 (*thrC*::*erm*, *amp*) (Guérout-Fleury et al. 1996).

**pSM59** (*amyE*::$P_{mdfA}$-*gfp kan*, *amp*) was designed to integrate a fusion of the promoter region of *mdfA* ($P_{mdfA}$) to a *gfp* reporter gene at the *B. subtilis amyE* locus. To construct this plasmid, a synthetic gene fragment containing the *mdfA* promoter, 5' leader sequence, RBS, and start codon, fused in-frame to *gfp* ($P_{mdfA}$-*gfp*), was assembled into EcoRI/BamHI-digested pER82 (*amyE*::*kan*, *amp*) (Driks et al. 1994). The *gfp* variant used in this construct harbored the *mut2* mutations (Cormack et al. 1996) as well as the monomerizing *A206K* mutation (Zacharias et al. 2002).

**pSM60** (*amyE*::$P_{mdfA}$-*gfp-mdfA kan*, *amp*) was designed to integrate a full-length *gfp-mdfA* fusion, under the control of the native *mdfA* promoter ($P_{mdfA}$), at the *B. subtilis amyE* locus. To construct this plasmid, a synthetic gene fragment containing the *mdfA* promoter, 5' leader sequence, RBS, and start codon, fused in-frame to *gfp* followed by a 23-amino acid linker (derived from pMMB759 (Berkmen and Grossman 2006)) and, finally, the full length *mdfA* coding sequence ($P_{mdfA}$-*gfp-mdfA*), was assembled into EcoRI/BamHI-digested pER82 (*amyE*::*kan*, *amp*) (Driks et al. 1994). The *gfp* variant used in this construct, as in pSM59, harbored the *mut2* mutations (Cormack et al. 1996) as well as the monomerizing *A206K* mutation (Zacharias et al. 2002).

**pSM44** (*αNTD-mdfA*), **pSM45** (*αNTD-mecA*), **pCS9** (*αNTD-mdfA*$^{E161K}$), **pSF9** (*αNTD-mdfA*$^{E53H}$), and **pXW12** (*αNTD-mdfA*$^{E116K}$) encode IPTG-inducible fusions of the N-terminal domain of the α subunit of RNA polymerase (αNTD) to MdfA, MecA, or MdfA variants, as indicated, for use as "prey" proteins in a transcription-based bacterial two-hybrid assay in *E. coli* (Dove and Hochschild 2004). To construct pSM44 and pSM45, the *mdfA* and *mecA* coding sequences were PCR amplified using primer pairs prSM89/prSM90 and prSM97/prSM98, respectively, and assembled into the NotI/BamHI-digested backbone of pBRα (Dove et al. 1997). To construct pCS9, pSF9, and pXW12, pSM44 was subject to site directed mutagenesis (Q5 Site-Directed Mutagenesis Kit, New England Biolabs) using primer pairs oCS3/oCS4 (*E161K*), oSF3/oSF4 (*E53H*), and oAK3/oAK4 (*E116K*), respectively.

**pSM49** (pACλcI-*clpC*), **pSM61** (pACλcI-*clpC*$^{Q11P}$), **pSM62** (pACλcI-*clpC*$^{N}$), **pSM63** (pACλcI-*clpC*$^{N,Q11P}$), **pSM67** (pACλcI-*clpC*$^{ND1M}$), **pSM70** (pACλcI-*clpC*$^{ND1M,Q11P}$), **pSM68** (pACλcI-*clpC*$^{D1M}$), and **pSM69** (pACλcI-*clpC*$^{D2}$), **pCS27** (pACλcI-*clpC*$^{ND1M,K12E}$), **pSF23** (pACλcI-*clpC*$^{ND1M,H79E}$), **pXW20** (pACλcI-*clpC*$^{ND1M,K85E}$) encode IPTG-inducible fusions of the bacteriophage λ cI protein (λcI) to the indicated variants of ClpC, for use as "bait" proteins in a transcription-based bacterial two-hybrid assay in *E. coli* (Dove and Hochschild 2004). To construct the plasmids harboring wild type *clpC* and *clpC*$^{Q11P}$ variants, the appropriate coding sequences were PCR amplified from PY79 chromosomal DNA (for wild type *clpC* variants) or CFB282 chromosomal DNA (for *clpC*$^{Q11P}$ variants) and assembled into the NotI/BamHI-digested backbone of pACλcI (Dove et al. 1997). The primer pairs used for PCR amplification were as follows: prSM87/prSM88 (*clpC* and *clpC*$^{Q11P}$), prSM127/prSM128 (*clpC*$^{N}$ and *clpC*$^{N,Q11P}$), prSM127/prSM151 (*clpC*$^{ND1M}$ and *clpC*$^{ND1M,Q11P}$), prSM152/prSM151 (*clpC*$^{D1M}$), and prSM153/prSM154 (*clpC*$^{D2}$). To construct the remaining plasmids, pSM67 was subject to site directed mutagenesis (Q5 Site-Directed Mutagenesis Kit, New England Biolabs) using primer pairs oCS8/oCS9 (*K12E*), oSF11/oSF12 (*H79E*), and oXW6/oXW7 (*K85E*).

**pET28a-*mdfA*** (*6xHis-thrombin-mdfA*) was designed to drive IPTG-inducible expression in *E. coli* of MdfA with an N-terminal 6xHis tag/thrombin cleavage site. To construct pET28a-*mdfA*, the full-length coding sequence of *mdfA* was PCR amplified with primers prSM35 and prSM36, and assembled into NheI/XhoI-digested pET28a (Novagen).

**pET151-*clpC*$^{N}$** (*6xHis-TEV-clpC*$^{N}$) and **pET151-*clpC*$^{N,Q11P}$** (*6xHis-TEV-clpC*$^{N,Q11P}$ in pET151) were designed to drive IPTG-inducible expression in *E. coli* of the ClpC N-terminal domain (residues 1-150; *WT* or harboring the *Q11P* substitution) with an N-terminal 6xHis tag/TEV cleavage site. First, a gene fragment encoding codon optimized $ClpC^{N}$ was synthesized and cloned into pET151 (Novagen), yielding pET151-*clpC*$^{N}$. This plasmid was then subjected to site-directed-mutagenesis (Q5 Site-Directed Mutagenesis Kit, New England Biolabs) using primers ClpC-Q11P-F and ClpC-Q11P-R, yielding pET151-*clpC*$^{N,Q11P}$.

**pCA528-*mdfA*** (*6xHis-SUMO-mdfA*) was designed to drive IPTG-inducible expression in *E. coli* of MdfA with an N-terminal 6xHis tag/SUMO cleavage site. The *mdfA* gene was amplified with primer pair IH236/IH237, digested with XhoI/BsmBI, and ligated into BsaI-digested pCA528 (Andréasson et al. 2008).

**pET28a-*mecA*** *(6xHis-thrombin-mecA)*, **pET28a-*clpC*** *(6xHis-thrombin-clpC)*, **pET28a-*clpC-DWB*** *(6xHis-thrombin-clpC*$^{E280A/E618A}$*)*, **pET28a-*clpC-Q11P*** (*6xHis-thrombin-clpC*$^{Q11P}$), **pET28a-*clpC-Q11A*** *(6xHis-thrombin-clpC*$^{Q11A}$*)*, and ***pET28a-clpP*** *(6xHis-thrombin-clpP)* were designed to drive IPTG-inducible expression in *E. coli* of MecA, ClpC, ClpC$^{DWB}$, ClpC$^{Q11P}$, ClpC$^{Q11A}$, or ClpP, respectively, with N-terminal 6xHis tags and thrombin cleavage sites. The respective genes were PCR amplified, digested with NcoI/XhoI, and ligated into NcoI/XhoI-digested pET28a (Novagen). The primers used for PCR amplification were as follows: IH220/IH221 *(mecA)*, IH17/IH27 *(clpC* and *clpC*$^{DWB}$*)*, IH50/IH27 *(clpC*$^{Q11P}$*)*, IH67/IH27 *(clpC*$^{Q11A}$*)*, and IH13/IH26 *(clpP)*. Note that DWB refers to the *E280A E618A* "double Walker B" *clpC* variant, the sequence of which was derived from Kirstein et al. (2006).

**Microscopy**

*B. subtilis* cells to be observed by microscopy were resuspended in phosphate-buffered saline (PBS) supplemented, when appropriate, with the membrane stain FM 4-64 (1 µg/mL, Molecular Probes) and/or the DNA stain 4',6-diamidino-2-phenylindole (DAPI; 1 µg/mL, Invitrogen). Cells were mounted on 1% agarose pads with poly-L-lysine treated coverslips and observed by phase and fluorescence microscopy with a Nikon Eclipse Ti-E inverted microscope equipped with a Nikon Plan Apochromat 100x phase objective and Chroma filter sets 49002 (excitation 470/40x, dichroic 495lpxr, and emission 525/50m), 49017 (excitation 560/40x, dichroic 590lpxr, and emission 590lp), and 49021 (excitation 405/20x, dichroic T425lpxr, and emission 460/50m) for detection of GFP, FM 4-64, and DAPI, respectively. Images were captured with an ORCA-Flash4.0 digital CMOS camera (Hamamatsu Photonics K.K.) using NIS-Elements Advanced Research software (Nikon Instruments Inc.). Images were pseudocolored, overlaid, and adjusted for brightness and contrast with Fiji software (Schindelin et al. 2012).

**Isolation and identification of *mdfA* extragenic suppressor mutants**

To isolate suppressor mutants immune to *mdfA*-mediated vegetative toxicity, a strain (SMB351) harboring two isopropyl-β-thiogalactopyranoside (IPTG)-inducible *mdfA* constructs, one at the *amyE* locus and the other at the *thrC* locus, was grown in LB to an optical density at 600 nm ($OD_{600}$) of approximately 0.6, and 100 µL was plated onto LB plates containing 1 mM IPTG. Plates were incubated overnight at 37°C and then at room temperature (25°C) for 3 additional days. Candidate suppressor colonies were restruck to LB plates containing 1 mM IPTG. Backcrosses into the wild type strain (PY79) were performed to identify and eliminate intragenic suppressor mutants located in one or both of the IPTG-inducible *mdfA* constructs. Suppressor mutants that did not demonstrate linkage to either IPTG-inducible *mdfA* construct were designated as extragenic suppressors. For these mutants, rare co-transformants (typically observed at a frequency of 1-10%) that displayed the suppression phenotype were selected for whole genome sequencing (Genewiz Inc.) to identify sites of mutations. Sequences were aligned and analyzed using Geneious software.

**Recombinant protein purification**

For the structural and biophysical studies of the interaction between MdfA and the ClpC N-domain ($ClpC^N$), T7 Express *lysY E. coli* cells (New England Biolabs) were transformed with plasmids containing full-length *mdfA* (pET28a-*yjbA*) or the N-terminal region of *clpC* (codons 1-150), either wild type (pET151-$clpC^N$) or the *Q11P* mutant (pET151-$clpC^{N,Q11P}$). Cells were grown in 0.5 L LB with appropriate antibiotics at 37°C to an $OD_{600}$ of 0.8 (for MdfA) or 0.5 (for $ClpC^N$ and $ClpC^{N,Q11P}$) at which point protein expression was induced with 0.5 mM IPTG for 16 h at 18°C (for MdfA) or for 4 h at 37°C (for $ClpC^N$ and $ClpC^{N,Q11P}$). For $^{15}N$-labeled $ClpC^N$ and $ClpC^{N,Q11P}$, growth and induction was carried out as described above but in M9 media supplemented with $^{15}N$-labeled ammonium chloride (Sigma-Aldrich) and $^{15}N$-ISOGRO (Sigma-Aldrich). Harvested cells were resuspended in lysis buffer (50 mM HEPES pH 7.5, 300 mM NaCl, 5 mM imidazole, 0.5 mM TCEP) supplemented with protease inhibitors (Roche cOmplete™ Mini EDTA-free Protease Inhibitor Cocktail, 1 mM PMSF), 10 µg/mL DNase, and 10 mM $MgCl_2$. Cells were lysed using a cell disruptor (Constant Systems Ltd.) and lysates were clarified by ultracentrifugation. Soluble 6xHis-tagged proteins were enriched by immobilized metal affinity chromatography (IMAC) on a 5 mL HisTrap™ HP column (GE Healthcare) and were eluted in buffer containing 300 mM imidazole, followed by dialysis into cleavage buffer (50 mM HEPES pH 7.5, 300 mM NaCl, 5 mM imidazole, 0.5 mM TCEP). $ClpC^N$ and $ClpC^{N,Q11P}$ proteins were digested with homemade TEV protease (100 µg/mL) at 4°C overnight to remove the 6xHis tag and were recovered in the flow-through of a second nickel IMAC purification. MdfA, $ClpC^N$, and $ClpC^{N,Q11P}$ proteins were subjected to size exclusion chromatography (SEC) with a HiLoad 16/60 Superdex 200 column (GE Healthcare) pre-equilibrated in buffers specific to downstream applications.

The full-length MdfA, MecA, $ClpC^{DWB}$, ClpC, $ClpC^{Q11P}$, $ClpC^{Q11A}$, and ClpP proteins used for ClpC oligomerization, ATPase, and protein degradation assays were purified from BL21(DE3) *E. coli* cells transformed with the appropriate plasmid (pCA528-*yjbA*, pET28a-*mecA*, pET28a-*clpC*-DWB, pET28a-*clpC*, pET28a-$clpC^{Q11P}$, pET28a-$clpC^{Q11A}$, and pET28a-*clpP* respectively. Cells were grown at 37°C to an $OD_{600}$ of 0.7-0.9 and expression was started by adding 1 mM IPTG. Cells were shifted to 22°C overnight for protein production. Purification of 6xHis-tagged proteins was performed by standard nickel affinity chromatography after cell lysis by French press in lysis buffer (50 mM Tris pH 8.0, 150 mM NaCl, 5 mM $MgCl_2$) (Turgay et al. 1997). Lysis buffer was supplemented with 20 mM or 250 mM imidazole during washing and elution steps, respectively. ClpC was purified as previously described (Turgay et al. 2001). Protein concentrations were calculated using the Bradford assay (Bio-Rad) and, if necessary, enriched using Vivaspin® spin columns (Sartorius). Proteins were frozen with liquid $N_2$ in aliquots in lysis buffer supplemented with 10% (w/v) glycerol and stored at -80°C.

**Analytical Size Exclusion Chromatography (SEC)**

For the analytical SEC experiments with MdfA and $ClpC^N$, the two proteins were purified as described in the “Recombinant protein purification” section above (final buffer conditions: 50 mM HEPES pH 7.5, 150 mM NaCl, 50 mM L-Glu, 0.5 mM TCEP) before being combined. MdfA, held constant at 266 μM, was run alone or with $ClpC^N$ at 1:0.5, 1:1 and 1:2 molar ratios, after incubation for 4 h at 4°C. The analytical SEC column (Superdex 75 Increase 10/300 GL, GE Healthcare) was pre-equilibrated in crystallization buffer (50 mM HEPES pH 7.5, 150 mM NaCl, 50 mM L-Glu, 0.5 mM TCEP). Due to the large difference in extinction coefficients, UV traces at 230 nm were recorded for analysis. The chromatography peaks were also analyzed by SDS-PAGE and Coomassie staining. Relevant fractions were used for subsequent crystallography experiments.

**Nuclear magnetic resonance (NMR)**

Unlabeled MdfA and $^{15}$N-labeled $ClpC^N$ and $ClpC^{N,Q11P}$ were purified as described in the “Recombinant protein purification” section above (final buffer conditions: 50 mM HEPES pH 7.5, 300 mM NaCl, 5 mM imidazole, 0.5 mM TCEP). 10% $D_2O$ (Sigma Aldrich) was added before protein was placed in 5 mm NMR tubes. 1D [$^1$H] and 2D [$^1$H,$^{15}$N]-HSQC spectra were recorded at 25°C using Bruker Avance NMR spectrometers, operating at 700 MHz and 800 MHz respectively with cryoprobes at the Biomolecular Spectroscopy Centre, King’s College London. 2D [$^1$H,$^{15}$N]-HSQC spectra were used to analyze the interaction between MdfA and $ClpC^N$ or $ClpC^{N,Q11P}$. Spectra were collected for $^{15}$N-labeled ClpC either alone or in the presence of unlabelled MdfA titrated up to 4-fold molar excess. Spectra were processed using NMRpipe (Delaglio et al. 1995) and analyzed using CCPNMR Analysis v.2 (Vranken et al. 2005).

A partial assignment of the $ClpC^N$ spectra was achieved by tracking peak movement during a pH titration as compared to $ClpC^N$ spectra deposited in the Biological Magnetic Resonance Bank (accession code 15383) (Kojetin et al. 2007). $ClpC^N$ was purified at pH 7.5 and the pH was adjusted to pH 6.6, 6, and 5.6 prior to [$^1$H,$^{15}$N]-HSQC spectra collection at 32°C.

**Protein Crystallography**

To form crystals of the MdfA/$ClpC^N$ complex, fractions from the gel filtration assay were concentrated (7-17 mg/mL), and set up in fine screens with hanging-drop vapor diffusion, with a 1 μL drop size, incubated at 16°C. Fine Screen 1 was based on the top apo MdfA hit from coarse screens (0.2 M $MgCl_2$, 0.1 M Tris pH 7.0, 10% [w/v] PEG 8000). Long needle crystals formed after 24 h, with the thickest crystals found at 17 mg/mL (in condition 0.2 M $MgCl_2$, 0.1 M Tris pH 7.0, 4% (w/v) PEG 8000) after 48 h. Fine Screen 2 was based on prior apo ClpC conditions (Wang et al. 2011) (2.0 M $(NH_4)_2SO_4$, 100 mM NaOAc, pH 4.6) and large hexagonal crystals formed after 48 h, with the best quality crystals found at 8.4 mg/mL. $ClpC^{N,Q11P}$ was screened at pH 8.85, 15 mg/ml, and best crystals formed in 200 mM $Li_2SO_4$, 100 mM NaOAc, pH 4.5, 50% w/v PEG 400.

All datasets were collected at Diamond Light Source. MdfA/$ClpC^N$ complex crystals on beamline I03 on a Pilatus 6M-F detector and a single wavelength of 0.9763 Å and 0.9795 Å respectively. $ClpC^{N,Q11P}$ data were collected on beamline I04-1 on a Eiger2 XE 16M with a single wavelength of 0.920 Å. Data were processed, merged and scaled in XIA2 3D (Winter et al.

2013) and XIA2 DIALS (Winter et al. 2018). Phaser (McCoy 2007) and MOLREP (Vagin and Teplyakov 2010) were used for molecular replacement, using PDB Accession 5HBN (Trentini et al. 2016) as a molecular replacement model. Data collection, processing and refinement statistics are shown in **Supplemental Table S6**.

To build the novel structure of MdfA, the molecular replacement solution was used as input to find the correct phases with the software SHELXE4MR (Thorn and Sheldrick 2013). Further automated model building with the SHELXE4MR solution was conducted with ARP/wARP Classic (Morris et al. 2003) and BUCCANEER_pipeline (Cowtan 2006), and manual rebuilding in Coot (Casañal et al. 2020). Refinement was performed in REFMAC5 (Murshudov et al. 2011) and PHENIX (Adams et al. 2010), with model validation and rebuilding using PDB_REDO (Joosten et al. 2014) and MolProbity (Williams et al. 2018) with reference to PDB Accession 3FES.

**Small-angle X-ray scattering (SAXS)**

SAXS experiments on isolated MdfA were performed at undulator beamline P12 at the PETRA III storage-ring35 at Deutsches Elektronen Synchrotron (DESY), Hamburg, Germany. The Pilatus 2M detector (DECTRIS, Switzerland) was located at a distance of 3.1 m from the sample. A freshly prepared BSA standard (3.0 mg/mL; 66 kDa) was measured for molecular weight determination after processing and normalization. Scattering data were collected at 0.3 mg/mL MdfA in buffer 50 mM HEPES, 50 mM L-glutamic acid, 150 mM NaCl, 0.5 mM TCEP, pH 8.85 at 25°C and a wavelength of $\lambda = 0.12$ nm, in batch mode (with 40 frames at an exposure time of 0.05 s per frame recorded). Data were analyzed with the ATSAS software suite v3.0.236.

### SUPPLEMENTAL TABLES

#### Table S1. *B. subtilis* strains used in this study

| Strain | Genotype[a,b] | Source or reference |
|---|---|---|
| PY79 | Prototrophic wild type | Youngman et al. 1984 |
| AHB324 | *ywrK*::Tn*917*::*amyE*::$P_{sspB}$-*lacZ cat* | Camp and Losick 2009 |
| AHB881 | *amyE*::$P_{spoIIQ}$-*lacZ cat* | Camp and Losick 2009 |
| AHB1296 | *amyE*::$P_{lonB}$-*lacZ cat* | This study |
| AHB1449 | *ylnF*::Tn*917*::*amyE*::$P_{spoIIQ}$-*T7RNAP spc*, *ywrK*::Tn*917*::*amyE*::$P_{T7}$-*lacZ cat* | Camp and Losick 2009 |
| AHB1560 | *ylnF*::Tn*917*::*amyE*::$P_{spoIIQ}$-*T7RNAP spc*, *ywrK*::Tn*917*::*amyE*::$P_{T7}$-*lacZ cat*, Δ*spoIIIAA-AH*::*erm*::*phleo* | This study |
| AHB1704 | *amyE*::$P_{yyaC}$-*lacZ cat* | This study |
| AHB1729 | *amyE*::$P_{csfC}$-*lacZ cat* | This study |
| AHB1841 | *ywrK*::Tn*917*::*amyE*::$P_{spoIIQ}$-*lacZ cat* | This study |
| AHB6207 | *amyE::*$P_{spoIIQ}$-*lacZ cat,* $clpC^{H79E}$ | This study |
| AHB6211 | *amyE::*$P_{spoIIQ}$-*lacZ cat,* $clpC^{K85E}$ | This study |
| AHB6213 | *amyE*::$P_{hyperspank}$-$mdfA^{E161K}$ *lacI spc*, $clpC^{K12E}$ | This study |
| AHB6215 | *amyE*::$P_{hyperspank}$-$mdfA^{E53H}$ *lacI spc*, $clpC^{H79E}$ | This study |
| AHB6220 | *amyE::*$P_{spoIIQ}$-*lacZ cat,* $yjbA^{E116K}$ | This study |
| AHB6229 | *amyE::*$P_{spoIIQ}$-*lacZ cat,* $clpC^{K85E}$, $yjbA^{E116K}$ | This study |
| AHB6231 | *amyE::*$P_{spoIIQ}$-*lacZ cat,* $clpC^{H79E}$, $yjbA^{E53H}$ | This study |
| AHB6234 | *amyE::*$P_{spoIIQ}$-*lacZ cat,* $clpC^{K12E}$, $yjbA^{E161K}$ | This study |
| AVB6 | *amyE::*$P_{spoIIQ}$-*lacZ cat,* $yjbA^{E53H}$ | This study |
| BAT87 | *amyE*::$P_{spoVT}$-*lacZ cat* | Gift of B. Traag |
| BZ184 | *amyE*::$P_{spoIID}$-*lacZ cat* | Margolis et al. 1993; gift of R. Losick |
| CFB123 | *amyE*::$P_{hyperspank}$-*lacZ lacI spc* | This study |
| CFB125 | *amyE*::$P_{hyperspank}$-[no *lacZ*] *lacI spc* | This study |
| CFB153 | *amyE*::$P_{spoIID}$-*lacZ cat*, Δ*mdfA*::*erm* | This study |
| CFB165 | *amyE*::$P_{spoIIP}$-*lacZ cat*, Δ*mdfA*::*erm* | This study |
| CFB189 | *amyE*::$P_{hyperspank}$-*mdfA lacI spc* | This study |
| CFB241 | *amyE*::$P_{hyperspank}$-*mdfA lacI spc*, Δ*clpC*::*erm* | This study |
| CFB270 | Δ*clpC*::*erm* | This study |
| CFB274 | *amyE*::$P_{hyperspank}$-*mdfA lacI spc*, Δ*clpP*::*erm* | This study |
| CFB282 | $clpC^{Q11P}$, Δ*yacL*::*erm* | This study |
| CFB302 | Δ*clpC*::*erm*, *amyE*::$P_{spoIIQ}$-*lacZ cat* | This study |
| CFB307 | *amyE*::$P_{spoIIQ}$-*lacZ cat,* $clpC^{Q11P}$, Δ*yacL*::*erm* | This study |
| CFB525 | *ywrK*::Tn*917*::*amyE*::$P_{xyl}$-*gfp-zapA cat* | This study |
| CFB530 | *ywrK*::Tn*917*::*amyE*::$P_{spoIIQ}$-*lacZ cat*, Δ*mdfA*::*erm* | This study |

| | | |
|---|---|---|
| CFB541 | *ywrK*::Tn*917*::*amyE*::P$_{spoIIQ}$-*lacZ cat*, Δ*mdfA*::*erm*, *amyE*::P$_{mdfA}$-*mdfA kan* | This study |
| CFB545 | *ywrK*::Tn*917*::*amyE*::P$_{spoIIQ}$-*lacZ cat*, Δ*mdfA*::*erm*, *amyE*::P$_{mdfA}$-*gfp-mdfA kan* | This study |
| CFB549 | *ywrK*::Tn*917*::*amyE*::P*xyl-GFP-zapA cat*, *amyE*::P$_{hyperspank}$-*mdfA lacI spc* | This study |
| CFB602 | Δ*mdfA*, *amyE*::P$_{hyperspank}$-*lacZ lacI spc* | This study |
| CFB604 | Δ*mdfA*, *amyE*::P$_{hyperspank}$-[no *lacZ*] *lacI spc* | This study |
| JDC138 | *amyE*::P1$_{csfB}$-*lacZ cat* | Flanagan et al. 2016 |
| CSB8 | *amyE*::P$_{hyperspank}$-*mdfA*$^{E161K}$ *lacI spc* | This study |
| PE511 | *amyE*::P$_{spoIIP}$-*lacZ cat* | Eichenberger et al. 2004; gift of R. Losick |
| RFB21 | *amyE*::P$_{hyperspank}$-*mdfA lacI spc*, *clpC*$^{K12E}$ | This study |
| RFB32 | *amyE*::*P*$_{spoIIQ}$-*lacZ cat, clpC*$^{K12E}$ | This study |
| SFB8 | *amyE*::P$_{hyperspank}$-*mdfA*$^{E53H}$ *lacI spc* | |
| SFB32 | *amyE*::P$_{hyperspank}$-*mdfA lacI spc*, *clpC*$^{Q11P}$ | This study |
| SFB38 | *amyE*::P$_{hyperspank}$-*mdfA lacI spc, clpC*$^{Q11A}$ | This study |
| SMB130 | *ylnF*::Tn*917*::*amyE*::P$_{spoIIQ}$-*T7RNAP spc*, *ywrK*::Tn*917*::*amyE*::P$_{T7}$-*lacZ cat*, Δ*spoIIIAA-AH*::*erm*::*phleo*, Δ*mdfA*::*erm* | This study |
| SMB165 | *ylnF*::Tn*917*::*amyE*::P$_{spoIIQ}$-*T7RNAP spc*, *ywrK*::Tn*917*::*amyE*::P$_{T7}$-*lacZ cat*, Δ*mdfA*::*erm* | This study |
| SMB179 | *ywrK*::Tn*917*::*amyE*::P$_{sspB}$-*lacZ cat*, Δ*mdfA*::*erm* | This study |
| SMB189 | *amyE*::P$_{spoIIQ}$-*lacZ cat*, Δ*mdfA*::*erm* | This study |
| SMB251 | Δ*mdfA*::*erm* | This study |
| SMB266 | *amyE*::P$_{mdfA}$-*lacZ cat* | This study |
| SMB296 | *amyE*::P$_{mdfA}$-*lacZ cat*, Δ*sigF*::*erm* | This study |
| SMB297 | *amyE*::P$_{mdfA}$-*lacZ cat*, Δ*sigE*::*erm* | This study |
| SMB298 | *amyE*::P$_{mdfA}$-*lacZ cat*, Δ*sigK*::*erm* | This study |
| SMB299 | *amyE*::P$_{mdfA}$-*lacZ cat*, Δ*sigG*::*kan* | This study |
| SMB300 | *amyE*::P$_{cotD}$-*lacZ cat* | This study |
| SMB301 | *amyE*::P$_{gerE}$-*lacZ cat* | This study |
| SMB302 | *amyE*::P1$_{csfB}$-*lacZ cat*, Δ*mdfA*::*erm* | This study |
| SMB303 | *amyE*::P$_{lonB}$-*lacZ cat*, Δ*mdfA*::*erm* | This study |
| SMB304 | *amyE*::P$_{yyaC}$-*lacZ cat*, Δ*mdfA*::*erm* | This study |
| SMB305 | *amyE*::P$_{csfC}$-*lacZ cat*, Δ*mdfA*::*erm* | This study |
| SMB310 | *amyE::P*$_{spoVT}$-*lacZ*, Δ*mdfA*::*erm* | This study |
| SMB332 | *amyE*::P$_{cotD}$-*lacZ cat*, Δ*mdfA*::*erm* | This study |
| SMB334 | *ywrK*::Tn*917*::*amyE*::P$_{sspB}$-*lacZ cat*, Δ*mdfA*::*erm*, *amyE*::P$_{mdfA}$-*mdfA kan* | This study |
| SMB335 | *amyE*::P$_{gerE}$-*lacZ cat*, Δ*mdfA*::*erm* | This study |

| | | |
|---|---|---|
| SMB352 | *amyE*::P$_{hyperspank}$-*mdfA lacI spc*, *clpC*$^{supp\#1}$<br>The *clpC*$^{supp\#1}$ allele harbors a +C frameshift insertion in the T251 codon (ACA>ACCA), causing truncation of the ClpC protein after K252 plus 5 incorrect amino acids. | This study |
| SMB355 | *amyE*::P$_{hyperspank}$-*mdfA lacI spc*, *clpC*$^{supp\#2}$<br>The *clpC*$^{supp\#2}$ allele harbors a -A frameshift deletion in the *clpC* K580 codon (AAA>AA), causing truncation of the ClpC protein after K580 plus 22 incorrect amino acids. | This study |
| SMB357 | *amyE*::P$_{hyperspank}$-*mdfA lacI spc*, *clpC*$^{supp\#5}$<br>The *clpC*$^{supp\#5}$ allele harbors a C>T nonsense mutation in the *clpC* Q676 codon (CAG>TAG), causing truncation of the ClpC protein after V675. | This study |
| SMB361 | *amyE*::P$_{hyperspank}$-*mdfA lacI spc*, *clpC*$^{supp\#7}$ (*clpC*$^{Q11P}$)<br>The *clpC*$^{supp\#7}$ allele harbors an A>C missense mutation in the *clpC* Q11 codon (CAA>CCA), switching the glutamine at position 11 of the ClpC protein to proline (Q11P). | This study |
| SMB365 | *amyE*::P$_{hyperspank}$-*mdfA lacI spc*, *clpC*$^{supp\#8}$<br>The *clpC*$^{supp\#8}$ allele harbors a 162 bp (54 codon) in-frame duplication of nucleotides 58-219 of the *clpC* coding sequence, causing the insertion of 54 amino acids between E73 and M74. | This study |
| SMB366 | *amyE*::P$_{hyperspank}$-*mdfA lacI spc*, *clpC*$^{supp\#12}$<br>The *clpC*$^{supp\#12}$ allele harbors a 18 bp (6 codon) in-frame duplication of nucleotides 1569-1587 of the *clpC* coding sequence, causing the insertion of 6 amino acids between A529 and G530. This allele also harbors a G>T missense mutation in the *clpC* E700 codon (GAG>GAT), switching the glutamate at position 700 of the ClpC protein to aspartate (E700D). | This study |
| SMB409 | *amyE*::P$_{mdfA}$-*gfp kan* | This study |
| SMB431 | Δ*mdfA* (in-frame) | This study |
| SMB434 | Δ*mdfA*::*erm*::*spc*, *amyE*::P$_{mdfA}$-*gfp-mdfA kan* | This study |
| SOB18 | *amyE*::P$_{hyperspank}$-*mdfA lacI spc*, *clpC*$^{H79E}$ | This study |
| SYB10 | *amyE*::*P*$_{spoIIQ}$-*lacZ cat, yjbA*$^{E161K}$ | This study |
| XWB12 | *amyE*::P$_{hyperspank}$-*mdfA*$^{E116K}$ *lacI spc* | This study |
| XWB50 | *amyE*::P$_{hyperspank}$-*mdfA lacI spc*, *clpC*$^{K85E}$ | This study |
| XWB58 | *amyE*::P$_{hyperspank}$-*mdfA*$^{E116K}$ *lacI spc*, *clpC*$^{K85E}$ | This study |

[a] All *B. subtilis* strains are isogenic with PY79 (Youngman et al. 1984).

[b] For information on the sources of gene deletions, *lacZ* fusions, and other constructs, see “Strain construction” section of Supplemental Materials and Methods.

**Table S2. Plasmids used in this study**

| **Plasmid** | **Description*** | **Source** |
|---|---|---|
| B. subtilis *loop-in/loop-out gene editing:* | | |
| pMiniMAD2 | *ori[ts] erm amp* | Patrick and Kearns 2008 |
| pSF14 | *clpC* in pMiniMAD2 | This study |
| pSF15 | *clpC*$^{Q11P}$ in pMiniMAD2 | This study |
| pSF19 | *clpC*$^{Q11A}$ in pMiniMAD2 | This study |
| pRF6 | *clpC*$^{K12E}$ in pMiniMAD2 | This study |
| pSO3 | *clpC*$^{H79E}$ in pMiniMAD2 | This study |
| pXW22 | *clpC*$^{K85E}$ in pMiniMAD2 | This study |
| B. subtilis *CRISPR-Cas9 gene editing:* | | |
| pAJS23 | *ori[ts] cas9 erm-gRNA kan* | Sachla et al. 2021 |
| pSY1 | *mdfA* repair template in pAJS23 | This study |
| pAV3 | *mdfA*$^{E53H}$ repair template in pAJS23 | This study |
| pRW3 | *mdfA*$^{E116K}$ repair template in pAJS23 | This study |
| pSY3 | *mdfA*$^{E161K}$ repair template in pAJS23 | This study |
| B. subtilis *integration/expression:* | | |
| pDR244 | *cre ori(ts) spc amp* | Meeske et al. 2015; Koo et al. 2017 |
| pEr::Sp | *erm::spc amp* | Steinmetz and Richter 1994 |
| pFG28 | *amyE*::P$_{xyl}$-*gfp-zapA cat amp* | Gueiros-Filho and Losick 2002 |
| pAH412 | *amyE*::P$_{cotD}$-*lacZ cat amp* | This study |
| pAH413 | *amyE*::P$_{gerE}$-*lacZ cat amp* | This study |
| pCS2 | *amyE*::P$_{hyperspank}$-*mdfA*$^{E161K}$ *lacI spc amp* | This study |
| pSF2 | *amyE*::P$_{hyperspank}$-*mdfA*$^{E53H}$ *lacI spc amp* | This study |
| pSM1 | *amyE*::P$_{mdfA}$-*mdfA kan amp* | This study |
| pSM12 | *amyE*::P$_{mdfA}$-*lacZ cat amp* | This study |
| pSM26 | *amyE*::P$_{hyperspank}$-*mdfA lacI spc amp* | This study |
| pSM30 | *thrC*::P$_{hyperspank}$-*mdfA lacI erm amp* | This study |
| pSM59 | *amyE*::P$_{mdfA}$-*gfp*$^{mut2,A206K}$ *kan amp* | This study |
| pSM60 | *amyE*::P$_{mdfA}$-*gfp*$^{mut2,A206K}$-*mdfA kan amp* | This study |
| pXW7 | *amyE*::P$_{hyperspank}$-*mdfA*$^{E116K}$ *lacI spc amp* | This study |
| E. coli *bacterial two-hybrid assay bait plasmids:* | | |
| pACλcI | *P*$_{lacUV5}$-directed synthesis of the λcI protein, plasmid marked by *cat* | Dove et al. 1997 |
| pSM49 | pACλcI-*clpC cat* | This study |
| pSM61 | pACλcI-*clpC*$^{Q11P}$ *cat* | This study |

| | | |
|---|---|---|
| pSM62 | pACλcI-*clpC*$^{N}$ *cat* | This study |
| pSM63 | pACλcI-*clpC*$^{N,Q11P}$ *cat* | This study |
| pSM67 | pACλCI-*clpC*$^{ND1M}$ *cat* | This study |
| pSM68 | pACλCI-*clpC*$^{D1M}$ *cat* | This study |
| pSM69 | pACλCI-*clpC*$^{D2}$ *cat* | This study |
| pSM70 | pACλCI-*clpC*$^{ND1M,Q11P}$ *cat* | This study |
| pCS27 | pACλCI-*clpC*$^{ND1M,K12E}$ *cat* | This study |
| pSF23 | pACλCI-*clpC*$^{ND1M,H79E}$ *cat* | This study |
| pXW20 | pACλCI-*clpC*$^{ND1M,K85E}$ *cat* | This study |
| E. coli *bacterial two-hybrid assay prey plasmids:* | | |
| pBRα | $P_{lacUV5}$/$P_{lpp}$-directed synthesis of the full length α subunit of *E. coli* RNAP, plasmid marked by *amp* | Dove et al. 1997 |
| pSM44 | pBRα-*mdfA amp* | This study |
| pSM45 | pBRα-*mecA amp* | This study |
| pCS9 | pBRα-*mdfA*$^{E161K}$ *amp* | This study |
| pSF9 | pBRα-*mdfA*$^{E53H}$ *amp* | This study |
| pXW12 | pBRα-*mdfA*$^{E116K}$ *amp* | This study |
| E. coli *protein expression:* | | |
| pET28a-*yjbA* (pSM27) | *6xHis-thrombin-mdfA* in pET28a | This study |
| pET151-*clpC*$^{N}$ | *6xHis-TEV-clpC*$^{N}$ (codon optimized) in pET151 | This study |
| pET151-*clpC*$^{N,Q11P}$ | *6xHis-TEV-clpC*$^{N,Q11P}$ (codon optimized) in pET151 | This study |
| pCA528-*mdfA* | *6xHis-SUMO-mdfA* in pCA528 | This study |
| pET28a-*mecA* | *6xHis-thrombin-mecA* in pET28a | This study |
| pET28a-*clpC*-DWB | *6xHis-thrombin-clpC*$^{E280A/E618A}$ in pET28a | This study |
| pET28a-*clpC* | *6xHis-thrombin-clpC* in pET28a | This study |
| pET28a-*clpC-Q11P* | *6xHis-thrombin-clpC*$^{Q11P}$ in pET28a | This study |
| pET28a-*clpC-Q11A* | *6xHis-thrombin-clpC*$^{Q11A}$ in pET28a | This study |
| *Also see "Plasmid construction" section of Supplemental Materials and Methods. | | |

**Table S3. Primers used in this study**

| Primer | Sequence (5’→3’)* | Description |
|---|---|---|
| AH62 | gatc**gaattc**ccttgcactatctcggagag | $P_{gerE}$ forward primer with EcoRI restriction site. Used to construct pAH413. |
| AH234 | gatgcggcatgcttatcagagaaag | $P_{cotD}$ forward primer. Used to construct pAH412. |
| AH383 | gatc**aagctt**catgataaaatcccctttccataaac | $P_{cotD}$ reverse primer with HindIII restriction site. Used to construct pAH412. |
| AH384 | gatc**aagctt**catgtattgtaaccctccttgctaaggtg | $P_{gerE}$ reverse primer with HindIII restriction site. Used to construct pAH413. |
| ClpC-Q11P-F | taccgaacgtgcacCgaaagttctggc | *clpC* (codon optimized) site-directed mutagenesis forward primer to generate *Q11P* mutation. Used to construct pET28a-*clpC-Q11P.* |
| ClpC-Q11P-R | gtgccagaactttcGgtgcacgttcgg | *clpC* (codon optimized) site-directed mutagenesis reverse primer to generate *Q11P* mutation. Used to construct pET28a-*clpC-Q11P.* |
| IH13 | ccg**ctcgag**ctttttgtcttctgtgtga | *clpP* reverse primer with XhoI restriction site. Used to construct pET28a-*clpP* |
| IH17 | catg**ccatgg**ggtttggaagatttacaga | *clpC* forward primer with NcoI restriction site. Used to construct pET28a-*clpC* and pET28a-*clpC-DWB*. |
| IH26 | catg**ccatgg**gtttaatacctacagtcattgaac | *clpP* forward primer with NcoI restriction site. Used to construct pET28a-*clpP* |
| IH27 | ccg**ctcgag**attcgttttagcagtcgtt | *clpC* reverse primer with XhoI restriction site. Used to construct pET28a-*clpC*, pET28a-*clpC-Q11P*, and pET28a-*clpC-Q11A*. |
| IH50 | catg**ccatgg**ggtttggaagatttacagaacgagctcCGa aagtactggc | *clpC*$^{Q11P}$ forward primer with NcoI restriction site. Used to construct pET28a-*clpC-Q11P*. |
| IH67 | catg**ccatgg**ggtttggaagatttacagaacgagctGCaa aagtactggc | *clpC*$^{Q11A}$ forward primer with NcoI restriction site. Used to construct pET28a-*clpC-Q11A*. |
| IH220 | tt**ccatgg**gccatcatcatcatcatcatcatgaaattgaaag aattaacgagcatac | *mecA* forward primer with NcoI restriction site. Used to construct pET28a-*mecA*. |
| IH221 | at**ctcgag**tcactatgatgcaaagtgttttttatcg | *mecA* reverse primer with XhoI restriction site. Used to construct pET28a-*mecA*. |

| | | |
|---|---|---|
| IH236 | tcga**ctcgag**TCAgttgacctttgattcgttttcc | *mdfA* reverse primer with XhoI restriction site and *mdfA* stop codon. Used to construct pCA528-*mdfA*. |
| IH237 | ccagtg**cgtctc**aggtggtATGttatttcttcatgatgtgtgg | *mdfA* forward primer with BsmBI restriction site and *mdfA* start codon. Used to construct pCA528-*mdfA*. |
| oAA2 | tactgtttcattacataCaTaacgatttgtcagagcttcc | *mdfA* site directed mutagenesis forward primer to generate *E53H* mutation. Used to construct pAV3. |
| oAA3 | tatgtaatgaaacagtactgacggcacccttaacag | *mdfA* reverse primer paired with oAA2 to generate *E53H* mutation. Used to construct pAV3. |
| oAK3 | cccaagacagAagcagctcgt | *mdfA* site directed mutagenesis forward primer to generate *E116K* mutation. Used to construct pXW7 and pXW12. |
| oAK4 | attagacggctttttctgacag | *mdfA* reverse primer paired with oAK3 to generate *E116K* mutation. Used to construct pXW7 and pXW12. |
| oCS3 | gacaagaaaaAaacggcaaattaag | *mdfA* site directed mutagenesis forward primer to generate *E161K* mutation. Used to construct pCS2 and pCS9. |
| oCS4 | agtcctctgacatgttctg | *mdfA* reverse primer paired with oCS3 to generate *E161K* mutation. Used to construct pCS2 and pCS9. |
| oCS8 | acgagctcaaGaagtactggcg | *clpC* site directed mutagenesis forward primer to generate *K12E* mutation. Used to construct pCS27 and pRF6. |
| oCS9 | tctgtaaatcttccaaacatc | *clpC* reverse primer paired with oCS8 to generate *H79E* mutation. Used to construct pCS27 and pRF6. |
| oRW4 | gtctaatcccaagacagAagcagctcgtttatgaaatgg | *mdfA* site directed mutagenesis forward primer to generate *E116K* mutation. Used to construct pRW3. |
| oRW5 | ctgtcttgggattagacggctttttctgacagggatc | *mdfA* reverse primer paired with oRW4 to generate *E116K* mutation. Used to construct pRW3. |
| oSF3 | tcattacataCaTaacgatttgtcagagcttcc | *mdfA* site directed mutagenesis forward primer to generate *E53H* mutation. Used to construct pSF2 and pSF9. |
| oSF4 | aacagtactgacggcacc | *mdfA* reverse primer paired with oSF3 to generate *E53H* mutation. Used to construct pSF2 and pSF9. |
| oSF5 | cctgcaggtcgactctaga**ggatcc**ggttccctacgcggtattc | *mcsB* forward primer. Used to construct pSF14. |

| | | |
|---|---|---|
| oSF6 | ttgtaaaacgacggccagt**gaattc**tctgctcaagctcttttaa gttag | *clpC* reverse primer. Used to construct pSF14. |
| oSF7 | agaacgagctcCaaaagtact | *clpC* site directed mutagenesis forward primer to generate *Q11P* mutation. Used to construct pSF15. |
| oSF8 | agaacgagctGCaaaagtactg | *clpC* site directed mutagenesis forward primer to generate *Q11A* mutation. Used to construct pSF19. |
| oSF9 | gtaaatcttccaaacatcatatc | *clpC* reverse primer paired with oSF7 or oSF8 to generate *Q11P* or *Q11A* mutation, respectively. Used to construct pSF15 and pSF19. |
| oSF11 | tcaaacgattGaAtatactcctagagctaaaaaagtcattg | *clpC* site directed mutagenesis forward primer to generate *H79E* mutation. Used to construct pSF23 and pSO3. |
| oSF12 | gacatttcctgcccgcgc | *clpC* reverse primer paired with oSF11 to generate *H79E* mutation. Used to construct pSF23 and pSO3. |
| oSY1 | **ggccaataaggcc**tttctagattaagaaataatc | pAJS23 forward primer. Used to construct pSY1. |
| oSY2 | **ggcctcgttggcc**gtcgac | pAJS23 reverse primer. Used to construct pSY1. |
| oSY3 | ctcactatagggtcgac**ggccaacgaggcc**gtcagagct gcaaacaactc | *mdfA* repair template forward primer. Used to construct pSY1. |
| oSY4 | atttcttaatctagaaa**ggccttattggcc**cagccaagcgg ctcagtg | *mdfA* repair template reverse primer. Used to construct pSY1. |
| oSY8 | gaggactgacaagaaaaAaacggcaaattaagcagctg | *mdfA* site directed mutagenesis forward primer to generate *E161K* mutation. Used to construct pSY3. |
| oSY9 | ttttcttgtcagtcctctgacatgttctggtgctaatg | *mdfA* reverse primer paired with oSY8 to generate *E116K* mutation. Used to construct pSY3. |
| oXW6 | tcctagagctGaaaaagtcattg | *clpC* site directed mutagenesis forward primer to generate *K85E* mutation. Used to construct pXW20 and pXW22. |
| oXW7 | gtataatgaatcgtttgagac | *clpC* reverse primer paired with oXW6 to generate *K85E* mutation. Used to construct pXW20 and pXW22. |
| prSM27 | gcggataacaattaagctta**gtcgac**taaggaggaactact atgttatttcttcatgatgtgtgg | *mdfA* forward primer with optimal RBS. Used to construct pSM26. |
| prSM28 | cctcgtttccaccgaattagctt**gcatgc**aagaaaaagacg ctaatcaaaaaacc | *mdfA* reverse primer. Used to construct pSM26. |
| prSM35 | tggtggtggtggtggtg**ctcgag**TCAgttgacctttgattcg | *mdfA* reverse primer including native stop codon. Used to construct pET28a-*mdfA*. |

| | | |
|---|---|---|
| prSM36 | tgccgcgcggcagccatatgATGttatttcttcatgatgtgtggg | *mdfA* forward primer including ATG start codon. Used to construct pET28a-*mdfA*. |
| prSM87 | gagacgtttggc**gcggccgc**aatgtttggaagatttacagaacg | *clpC* forward primer. Used to construct pSM49, pSM61. |
| prSM88 | gatctgtaaggtaa**ggatcc**TTAattcgttttagcagtcg | *clpC* reverse primer including native stop codon. Used to construct pSM49, pSM61. |
| prSM89 | gagaaaccagag**gcggccgc**attatttcttcatgatgtgtggg | *mdfA* forward primer. Used to construct pSM44. |
| prSM90 | gcgtccggcgtaga**ggatcc**tcagttgacctttgaattcgttttcc | *mdfA* reverse primer. Used to construct pSM44. |
| prSM97 | gagaaaccagag**gcggccgc**agaaattgaaagaattaacgagc | *mecA* forward primer. Used to construct pSM45. |
| prSM98 | gcgtccggcgtaga**ggatcc**ctatgatgcaaagtgtttttttatcg | *mecA* reverse primer. Used to construct pSM45. |
| prSM127 | gagacgtttggc**gcggccgc**aatgtttggaagatttacagaacgagc | *clpC* forward primer. Used to construct pSM62, pSM63, pSM67, and pSM70. |
| prSM128 | gatctgtaaggtaa**ggatcc**TTAattacttcctagaagctggagc | *clpC* reverse primer with engineered stop codon. Used to construct pSM62 and pSM63. |
| prSM151 | gatctgtaaggtaa**ggatcc**TTAggtccagctggatacaaccatcgc | *clpC* reverse primer with engineered stop codon. Used to construct pSM67, pSM68, and pSM70. |
| prSM152 | gagacttttggc**gcggccgc**atcagcggcaggaacaaacagcaatgcg | *clpC* forward primer. Used to construct pSM68. |
| prSM153 | gagacgtttggc**gcggccgc**aaaaatcgcccaaactgaaactgataagc | *clpC* forward primer. Used to construct pSM69. |
| prSM154 | gatctgtaaggtaa**ggatcc**TTAattcgttttagcagtcgttttacg | *clpC* reverse primer including native stop codon. Used to construct pSM69. |
| pr*yjbA*FC | tggtagcgaccggcgctca**ggatcc**ctttttatgattcttacataaaccc | $P_{mdfA}$ forward primer. Used to construct pSM1. |
| pr*yjbA*PF | gctgtcaaacatga**gaattc**ctttttatgattcttacataaacccagtgattcc | $P_{mdfA}$ forward primer. Used to construct pSM12. |
| pr*yjbA*PR | cgtcagtaacttccac**aagctt**catctctaacccctcacttttataaaaactg | $P_{mdfA}$ reverse primer. Used to construct pSM12. |
| pr*yjbA*RC | ggggtaccgagctc**gaattc**tcagttgacctttgattcg | *mdfA* reverse primer. Used to construct pSM1. |

*Relevant restriction endonuclease recognition sites are in bold. Relevant substitutions, insertions, or features are indicated in uppercase. Sequences that provide complementarity for assembly are underlined.

### Table S4. Gene fragments used in this study

| gBlock / Description | Sequence (5'→3')* |
|---|---|
| $P_{mdfA}$-*gfp* / *gfp*$^{mut2,A206K}$ fused to the *mdfA* promoter. Used to construct pSM59. | ggtaatggtagcgaccggcgctca**ggatcc**ctttttatgattcttacataaacccagtgattccagatttcctcaa<br>tctctttggaataatttctctatcccttgcctatactgtggacaacagtttttataaaagtgaggggttagagatgaa<br>aggagaagaacttttcactggagttgtcccaattcttgttgaattagatggtgatgttaatgggcacaaattttctgt<br>cagtggagagggtgaaggtgatgcaacatacggaaaacttacccttaaatttatttgcactactggaaaacta<br>cctgttccatggccaacacttgtcactactttcgcgtatggtcttcaatgctttgcgagatacccagatcatatgaa<br>acagcatgactttttcaagagtgccatgcccgaaggttatgtacaggaaagaactatatttttcaaagatgacg<br>ggaactacaagacacgtgctgaagtcaagtttgaaggtgatacccttgttaatagaatcgagttaaaaggtatt<br>gattttaaagaagatggaaacattcttggacacaaattggaatacaactataactcacacaatgtatacatcat<br>ggcagacaaacaaaagaatggaatcaaagttaacttcaaaattagacacaacattgaagatggaagcgttc<br>aactagcagaccattatcaacaaaatactccaattggcgatggccctgtccttttaccagacaaccattacctgt<br>ccacacaatctaagctttcgaaagatcccaacgaaaagagagaccacatggtccttcttgagtttgtaacagc<br>tgctgggattacacatggcatggatgaactatacaaataa**gaattc**gagctcggtacccctggatttcac |
| $P_{mdfA}$-*gfp-mdfA* / Full-length *gfp*$^{mut2,A206K}$-*mdfA* fusion gene, fused to the *mdfA* promoter. Used to construct pSM60. | ggtaatggtagcgaccggcgctca**ggatcc**ctttttatgattcttacataaacccagtgattccagatttcctcaa<br>tctctttggaataatttctctatcccttgcctatactgtggacaacagtttttataaaagtgaggggttagagatgaa<br>aggagaagaacttttcactggagttgtcccaattcttgttgaattagatggtgatgttaatgggcacaaattttctgt<br>cagtggagagggtgaaggtgatgcaacatacggaaaacttacccttaaatttatttgcactactggaaaacta<br>cctgttccatggccaacacttgtcactactttcgcgtatggtcttcaatgctttgcgagatacccagatcatatgaa<br>acagcatgactttttcaagagtgccatgcccgaaggttatgtacaggaaagaactatatttttcaaagatgacg<br>ggaactacaagacacgtgctgaagtcaagtttgaaggtgatacccttgttaatagaatcgagttaaaaggtatt<br>gattttaaagaagatggaaacattcttggacacaaattggaatacaactataactcacacaatgtatacatcat<br>ggcagacaaacaaaagaatggaatcaaagttaacttcaaaattagacacaacattgaagatggaagcgttc<br>aactagcagaccattatcaacaaaatactccaattggcgatggccctgtccttttaccagacaaccattacctgt<br>ccacacaatctaagctttcgaaagatcccaacgaaaagagagaccacatggtccttcttgagtttgtaacagc<br>tgctgggattacacatggcatggatgaactatacaaaggagaaggacaaggacaaggacaaggaccagg<br>acgaggatacgcataccgaagcggttccggaatgttatttcttcatgatgtgtgggtcaattggtttgaagggga<br>agaaaatggctacaatgtatgccactttcacgaatggcgcaaggaagatacggttgagcttttggatcaggtg<br>ccgctgttaagggtgccgtcagtactgtttcattacatagaaaacgatttgtcagagcttccgaaaggtttgcttg<br>aagacgtacatcagaaatcgtacattcgaaaaaatcatgagcggacaaagcttgagtattgttttgtcgtaaca<br>gacggaatcggcattttagctgttgacacgatcggctacacgatccctgtcagaaaaagccgtctaatcccaa<br>gacaggagcagctcgtttatgaaatggtaaaagatgtagagcctgaaacatatgaatttgagccaaagaag<br>ctggaatcctccaaggaatatcacatcttatcattagcaccagaacatgtcagaggactgacaagaaaagaa<br>cggcaaattaagcagctgatgtttatggcgcttgaccaattaaaagggctgaaaaaccgagcggaaattggc<br>tactggtacacagagtggaatccgcatatgtatgaacaaatcaaaaggatgagctttgaagagatttgggac<br>atgctgtataacgaaacaattgaaggctggtcggacaaacacctagctttttgcgaaaacctaataaaagga<br>cagcctttttcgaaaagctttgggaaatggaaaacgaatcaaaggtcaactga**gaattc**gagctcggtaccc<br>ctggatttcac |

*Sequences that provide complementarity for isothermal assembly are underlined. Relevant substitutions or insertions are indicated in uppercase.

**Table S5. Known and putative $\sigma^F$ regulon genes included in the metabolic differentiation factor candidate screen**

| Gene[a] | Knockout[b] | Test strain[c] | Evidence for inclusion in $\sigma^F$ regulon |
|---|---|---|---|
| *csfB (gin)* | BKE00240 | SMB120 | Arrieta-Ortiz et al. 2015; Decatur and Losick 1996; Flanagan et al. 2016; Serrano et al. 2011; Wang et al. 2006 |
| *rnmV* | BKE00410 | SMB121 | In an operon with *ksgA* |
| *ksgA* | BKE00420 | SMB131 | Wang et al. 2006 |
| *fin (yabK)* | BKE00540 | SMB122 | Arrieta-Ortiz et al. 2015; Camp et al. 2011; Steil et al. 2005 |
| *mfd* | BKE00550 | SMB123 | Possibly in an operon with *fin* |
| *yabT* | BKE00660 | SMB124 | Arrieta-Ortiz et al. 2015; Wang et al. 2006 |
| *yetF* | BKE07140 | SMB47 | Arrieta-Ortiz et al. 2015; Steil et al. 2005 |
| *hypO (yfkO)* | BKE07830 | SMB48 | Steil et al. 2005 |
| *yfhD* | BKE08490 | SMB125 | Arrieta-Ortiz et al. 2015; Kuwana et al. 2002; Steil et al. 2005 |
| *yfhE* | BKE08500 | SMB126 | Arrieta-Ortiz et al. 2015; Steil et al. 2005 |
| *yfhF* | BKE08510 | SMB127 | In an operon with *yfhD* and *yfhE* |
| *queG (yhbA)* | BKE08910 | SMB49 | Steil et al. 2005 |
| *yhcM* | BKE09140 | SMB50 | Arrieta-Ortiz et al. 2015; Kuwana et al. 2002; Steil et al. 2005 |
| *yhcN* | BKE09150 | SMB128 | Bagyan et al. 1998b; Wang et al. 2006 |
| *yhfM* | BKE10280 | SMB51 | Arrieta-Ortiz et al. 2015; Steil et al. 2005; Wang et al. 2006 |
| *yhfW* | BKE10390 | SMB52 | Arrieta-Ortiz et al. 2015; Steil et al. 2005; Wang et al. 2006 |
| *yisN* | BKE10780 | SMB129 | Steil et al. 2005 |
| *mdfA (yjbA)* | BKE11410 | SMB130 | Arrieta-Ortiz et al. 2015; Steil et al. 2005; Wang et al. 2006 |
| *ykuS* | BKE14200 | SMB53 | Arrieta-Ortiz et al. 2015; Steil et al. 2005 |
| *ylbB* | BKE14950 | SMB147 | Arrieta-Ortiz et al. 2015; Steil et al. 2005; Wang et al. 2006 |
| *ylbC* | BKE14960 | SMB148 | Arrieta-Ortiz et al. 2015; Steil et al. 2005; Wang et al. 2006 |
| *miaA* | BKE17330 | SMB150 | Arrieta-Ortiz et al. 2015; Steil et al. 2005 |
| *sspN* | BKE18020 | SMB151 | Arrieta-Ortiz et al. 2015; Steil et al. 2005; Wang et al. 2006 |
| *tlp* | BKE18030 | SMB190 | Arrieta-Ortiz et al. 2015; Steil et al. 2005 |
| *ypzH* | BKE22849 | SMB191 | Arrieta-Ortiz et al. 2015 |
| *seaA* | BKE22850 | SMB192 | Arrieta-Ortiz et al. 2015; Steil et al. 2005 |
| *yphA* | BKE22860 | SMB113 | Arrieta-Ortiz et al. 2015; Decatur and Losick 1996; Steil et al. 2005 |
| *dacF* | BKE23480 | SMB112 | Arrieta-Ortiz et al. 2015; Schuch and Piggot 1994; Steil et al. 2005; Wang et al. 2006 |
| *ripX* | BKE23510 | SMB54 | Steil et al. 2005 |
| *spoIVB* | BKE24230 | SMB116 | Arrieta-Ortiz et al. 2015; Gomez and Cutting 1996; Steil et al. 2005; Wang et al. 2006 |
| *yqhQ* | BKE24490 | SMB195 | Arrieta-Ortiz et al. 2015; Wang et al. 2006 |
| *yqhP* | BKE24500 | SMB238 | Arrieta-Ortiz et al. 2015 |
| *yqhH* | BKE24580 | SMB196 | Arrieta-Ortiz et al. 2015; Steil et al. 2005; Wang et al. 2006 |

| | | | |
|---|---|---|---|
| *yqhG* | BKE24590 | SMB197 | Arrieta-Ortiz et al. 2015; Steil et al. 2005; Wang et al. 2006 |
| *yqzG* | BKE24650 | SMB55 | Arrieta-Ortiz et al. 2015; Steil et al. 2005; Wang et al. 2006 |
| *gpr* | BKE25540 | SMB198 | Arrieta-Ortiz et al. 2015; Steil et al. 2005; Sussman and Setlow 1991; Wang et al. 2006 |
| *yrrS* | BKE27300 | SMB199 | Steil et al. 2005 |
| *pbpI (yrrR)* | BKE27310 | SMB200 | Arrieta-Ortiz et al. 2015; Steil et al. 2005; Wang et al. 2006 |
| *yrzK* | BKE27570 | SMB201 | Arrieta-Ortiz et al. 2015; Steil et al. 2005 |
| *bofC* | BKE27750 | SMB202 | Arrieta-Ortiz et al. 2015; Gomez and Cutting 1997; Steil et al. 2005; Wang et al. 2006 |
| *lonB* | BKE28210 | SMB204 | Arrieta-Ortiz et al. 2015; Serrano et al. 2001; Steil et al. 2005; Wang et al. 2006 |
| *gerW (ytfJ)* | BKE29500 | SMB133 | Arrieta-Ortiz et al. 2015; Kuwana et al. 2002; Steil et al. 2005; Wang et al. 2006 |
| *ytfI* | BKE29510 | SMB134 | Arrieta-Ortiz et al. 2015; Steil et al. 2005; Wang et al. 2006 |
| *rppH (mutTA)* | BKE30630 | SMB135 | Ramírez et al. 2004; Wang et al. 2006 |
| *yuiC* | BKE32070 | SMB136 | Arrieta-Ortiz et al. 2015; Steil et al. 2005; Wang et al. 2006 |
| *yutH* | BKE32270 | SMB137 | Steil et al. 2005 |
| *yusW* | BKE32950 | SMB56 | Arrieta-Ortiz et al. 2015; Steil et al. 2005 |
| *gerAA* | BKE33050 | SMB138 | Wang et al. 2006 |
| *gerAB* | BKE33060 | SMB139 | Wang et al. 2006 |
| *gerAC* | BKE33070 | SMB140 | Wang et al. 2006 |
| *ywnJ* | BKE36540 | SMB141 | Arrieta-Ortiz et al. 2015; Steil et al. 2005; Wang et al. 2006 |
| *spoIIQ* | BKE36550 | SMB117 | Arrieta-Ortiz et al. 2015; Londoño-Vallejo et al. 1997; Steil et al. 2005; Wang et al. 2006 |
| *ywnF* | BKE36580 | SMB143 | Steil et al. 2005 |
| *ywlB* | BKE36960 | SMB142 | Arrieta-Ortiz et al. 2015; Steil et al. 2005 |
| *spoIIR* | BKE36970 | SMB118 | Arrieta-Ortiz et al. 2015; Karow et al. 1995; Steil et al. 2005; Wang et al. 2006 |
| *pbpG (ywhE)* | BKE37510 | SMB119 | Arrieta-Ortiz et al. 2015; Pedersen et al. 2000; Wang et al. 2006 |
| *rsfA* | BKE37620 | SMB144 | Arrieta-Ortiz et al. 2015; Steil et al. 2005; Wang et al. 2006; Wu and Errington 2000 |
| *katX* | BKE38630 | SMB145 | Arrieta-Ortiz et al. 2015; Bagyan et al. 1998a; Petersohn et al. 1999; Wang et al. 2006 |
| *yyaC* | BKE40950 | SMB146 | Arrieta-Ortiz et al. 2015; Steil et al. 2005; Wang et al. 2006 |

[a] Genes are listed in order of their position on the *B. subtilis* chromosome. Note that not every gene known or predicted to be activated by $\sigma^F$ was included in the screen.

[b] Erythromycin-marked gene deletions were from the *Bacillus subtilis* “BKE” Knockout Collection (Koo et al. 2017), distributed by the *Bacillus* Genetic Stock Center.

[c] Test strains were constructed by transforming strain AHB1560 (see **Supplemental Table S1**) with chromosomal DNA derived from the relevant BKE deletion strain.

**Table S6. Data collection, processing and refinement statistics for MdfA/ClpC$^{N}$ and ClpC$^{N,Q11P}$ crystal structures.**

| **Protein** | **MdfA/ClpC$^{N}$** | **ClpC$^{N,Q11P}$** |
|---|---|---|
| Beamline | Diamond Light Source I03 | Diamond Light Source I04-1 |
| Wavelength | 0.9763 Å | 0.9159 Å |
| Beam size | 50 x 20 µm | 60 x 50 µm |
| Transmission | 50.16 % | 47.00 % |
| Flux | 5.13 E+11 | 1.65 E+11 |
| Exposure time | 0.013s | 0.100 s |
| Number of images | 5960 | 3600 |
| Ω Osc | 0.05° | 0.10° |
| Data processing | xia2 3d | xia2 dials |
| Resolution Range | 2.02 - 29.90 (2.02 - 2.05) Å | 1.15 - 42.31 (1.15 - 1.17) Å |
| Space Group | P 31 | P 65 |
| Unit Cell | 69.82, 69.82,<br>89.71, 90.00,<br>90.00, 120.00 | 84.62, 84.62,<br>32.16, 90.00,<br>90.00, 120.00 |
| Total Reflections | 276606 (14426) | 894902 (36935) |
| Unique Reflections | 32111 (1644) | 45932 (2233) |
| Multiplicity | 8.6 (8.8) | 19.5 (16.5) |
| Completeness | 100 (100) % | 97.7 (96.3) % |
| Mean I/Sigma(I) | 9.4 (0.9) | 14.9 (1.3) |
| Wilson B-factor | 35.066 | 14.657 |
| R-meas | 0.149 (2.153) | 0.098 (12.009) |
| CC1/2 | 1.0 (0.5) | 1.0 (0.7) |
| Reflections used in refinement | 27283 | 43587 |
| Reflections used for $R_{free}$ | 1395 | 2288 |
| Final $R_{work}$ | 0.176 | 0.145 |
| Final $R_{free}$ | 0.234 | 0.166 |
